# Endosomal spatio-temporal modulation of the cortical RhoA zone conditions epithelial cell organization

**DOI:** 10.1101/2020.04.19.044891

**Authors:** Gaston Cécile, De Beco Simon, Doss Bryant, Pan Meng, Gauquelin Estelle, D’Alessandro Joseph, Lim Chwee Teck, Ladoux Benoit, Delacour Delphine

**Affiliations:** Cell adhesion and mechanics, Institut Jacques Monod, CNRS UMR7592, Paris Diderot University, 75205 Paris Cedex 13, France; Mechanobiology Institute, T-lab, Singapore 117411, Singapore

**Keywords:** cell plasticity, polarity, contractility, RhoA, cell migration

## Abstract

At the basis of cell shape and behavior, actomyosin organization and force-generating property are widely studied, however very little is known about the regulation of the contractile network in space and time. Here we study the role of the epithelial-specific protein EpCAM, a contractility modulator, in cell shape and motility, and we show that it is required for the maturation of stress fibers and frontrear polarity acquisition at the single cell level. There, EpCAM ensures the remodeling of a transient active RhoA zone in the cortex of spreading epithelial cells. GTP-RhoA follows the endosomal pathway mediated by Rab35 and EHD1, where it co-evolves together with EpCAM. In fact, EpCAM balances GTP-RhoA turnover in order to tune actomyosin remodeling for cell shape, polarity and mechanical property acquisition. Impairment of GTP-RhoA endosomal trafficking either by EpCAM silencing or Rab35 / EHD1 mutant expression prevents correct myosin-II activity, stress fiber formation, and ultimately cell polarization. Collectively, this work shows that the coupling of EpCAM/RhoA co-trafficking to actomyosin rearrangement is critical for spreading, and advances our understanding of how biochemical and mechanical properties can be coupled for cell plasticity.

## Introduction

Biological processes as diverse as cell division, extrusion, maintenance of cell shape or morphogenetic movements rely on the proper regulation of the contractile network patterning and activity (Rosenblatt et al., 2001; Yamada and Nelson, 2007; Abe and Takeichi, 2008; Papusheva and Heisenberg, 2010; Green et al., 2012; Munjal and Lecuit, 2014; Munjal et al., 2015; Heer and Martin, 2017; Hannezo and Heisenberg, 2019). Biochemical and mechanical inputs’ coupling is increasingly studied (Hannezo and Heisenberg, 2019) and, as places of actin remodeling and force generation and transmission, stress fibers are privileged sites to tackle this issue (Vogel and Sheetz, 2006; Ladoux et al., 2016).

Stress fibers are formed by cross-linked actin bundles that display an alternate pattern of α-actinin and myosin-IIA, are attached at both ends to FAs, and constitute the force generating fibers (Burridge and Wittchen, 2013). Radial fibers, devoid of myosin-IIA perpendicularly elongate from focal adhesion at cell periphery and connect circumferential actin arcs which arise from the bundling of short actin filaments following the actin retrograde flow near the dorsal surface of the cell (Hotulainen and Lappalainen, 2006; Tojkander et al., 2012). Circumferential arcs also exert contractile forces that are collectively transmitted to radial fibers, which in turn passively transmit tension from the cell center to the FA anchoring them (Burnette et al., 2014). As such, radial fibers exert low forces on the substrate (Soiné et al., 2015; Lee et al., 2018). A first study by Hotulainen and colleagues demonstrated that radial fibers grow by formin-mediated actin polymerization at FAs, and that ventral stress fibers arise from the fusion of radial fibers and associated-transverse arcs following the arcs contraction during the retrograde flow of the SF (Hotulainen and Lappalainen, 2006; Tojkander et al., 2015; Tee et al., 2015). Another study reported the existence of stress fiber assembly *de novo* from the concatenation of short actin filaments (Machesky and Hall, 1997; Vallenius, 2013). Stress fiber formation and dynamics have been actively scrutinized in fibroblasts and mesenchymal cells, where symmetry breaking and acquisition of front-rear polarity, as well as stress fiber maturation requires α4-actinin and contractility (Hotulainen and Lappalainen, 2006; Prager-Khoutorsky et al., 2011; Roca-Cusachs et al., 2013; Gupta et al., 2015).

Force generation and regulation rely on the presence and the activity of the non-muscle myosin-II motor. Phosphorylation of its regulatory light chain (MRLC) on serine 19 (S19) and threonine (T18) favors myosin-II ATPase activity and its assembly with actin filaments (Craig et al., 1983; Watanabe et al., 2007; Vicente-Manzanares and Horwitz, 2010). In addition, myosin-II dephosphorylation is required for the motor displacement along actin fibers (Watanabe et al., 2007). Several signaling pathways modulate myosin-II activity, among which the Rho pathway has been extensively studied (Hall, 1998; Spiering and Hodgson, 2011; Hodge and Ridley, 2016). In fact, Rho-associated coiled coil-containing kinases (ROCK) promote contractility by MRLC phosphorylation but also by inactivating the Myosin Light Chain Phosphatase (MLCP) (Amano et al., 2000; Vicente-Manzanares et al., 2009). Upstream, the small GTPase RhoA mediates ROCK stimulation (Julian and Olson, 2014), therefore its distribution and activity levels must be tightly controlled to ensure the correct contractile response within the cell (Agarwal and Zaidel-Bar, 2019). RhoGTPases are described as molecular switches, as they transition between an activated (GTP-loaded) and inactivated (GDP-loaded) state. This cycling between inactive and active form is important for the maintenance of active RhoA zones, and would occur near the plasma membrane (Rossman et al., 2005; Bos et al., 2007; Hodge and Ridley, 2016). Although RhoA activation facilitates its translocation to the plasma membrane (Michaelson et al., 2001), it is now clear that RhoA inactivation is as important as its activation, since it enables a pulsatile behavior of myosin-II activity needed for efficient contractility (Mason et al., 2016; Teo and Yap, 2016). Current research efforts particularly focus on understanding the upstream controlling mechanisms of this canonical contractility regulator.

Being subjected to the most acute remodeling events, epithelial tissues are especially sensitive to any cues affecting their tensional homeostasis and have thus received a lot of attention in the study of this process. EpCAM (<u>E</u>pithelial <u>C</u>ell <u>A</u>dhesion <u>M</u>olecule) is a transmembrane protein, exclusively expressed in epithelial cells in physiological conditions, and primarily described as a Ca2+-independent cell-cell adhesion molecule crucial for epithelial integrity (Litvinov et al., 1994; Balzar et al., 1998). During early stages of zebrafish or *Xenopus* development, EpCAM’s extinction leads to epiboly defects and numerous lesions of the future epidermis (Slanchev et al., 2009; Maghzal et al., 2013). In addition, the loss of EpCAM leads to the development of a rare human disease so-called Congenital Tufting Enteropathy (CTE), characterized by the formation of distinctive lesions in the intestinal epithelium (Patey et al., 1997; Salomon et al., 2014, 2017). However, the conflicting functional interaction of EpCAM with E-cadherin and recent structural analyses tend to challenge its initial function (Litvinov et al., 1997; Winter et al., 2003, 2007; Pavšič et al., 2014; Gaber et al., 2018), and emphasize the need to reconsider its mechanism of action. A link between EpCAM and the actin cytoskeleton was proposed in the 1990’s, with an influence of EpCAM deprivation or overexpression on actin organization (Guillemot et al., 2001), but how it would influence the actin network remained unclear. More recently, several reports highlighted EpCAM as an intriguing regulator of actomyosin contractility for the organization of epithelial assemblies. In *Xenopus* ectoderm explants, Fagotto and colleagues demonstrated an internalization of C-Cadherin and an increase of cell contractility under EpCAM’s deprivation, which was further confirmed in human Caco2 cells (Maghzal et al., 2010, 2013). Moreover, EpCAM’s silencing triggers an inappropriate distribution and magnitude of actomyosin activity at tricellular contacts, impacting the epithelial apico-basal polarity and the global monolayer arrangement in CTE patients and Caco2 cells (Salomon et al., 2017; Gaston et al., 2017). The impact of EpCAM on cell contractility modulation was pioneered by Fagotto and colleagues, who showed that the excess of cell contractility in mutant *Xenopus* explants was under the control of nPKC-dependent Erk signaling (Maghzal et al., 2010, 2013). However, the Erk signaling pathway is likely not the only mechanism involved. In fact, within the same studies, dominant negative (dnRhoA) expression or drug treatments affecting RhoA signaling were equally effective in restoring a normal phenotype in EpCAM-MO explants than PKCη inhibitor treatment. Although dnRhoA expression was even more effective in limiting the loss of integrity in EpCAM-MO explants than treatment with a PKC-negative dominant (Maghzal et al., 2013), the participation of the RhoA pathway in the EpCAM’s mechanism of action was not further pursued by the authors. Given these results and the numerous feedback controls between the different pathways regulating contractility, the molecular mechanism linking EpCAM to cellular contractility deserves therefore further analyses.

Here we aimed to understand the participation of EpCAM in cell polarization and actomyosin organization. Facing the diversity and complexity of interconnected regulating mechanisms in monolayers, we decided to conduct this study in isolated cells. We reveal that EpCAM plays a role independently of cell-cell contacts in single cell polarization. There, it controls the development of proper contractility for stress fiber maturation and self-organization through RhoA signaling modulation. From a mechanistic point of view, we show that EpCAM is required for the endosomal remodeling of an active RhoA zone at the cell cortex during cell spreading and polarization.

## Results

### EpCAM is required for cell polarization independently of cell-cell contacts

In an epithelial context, EpCAM has been exclusively studied in cell clusters and monolayers (Schnell et al., 2013; Mueller et al., 2014; Herreros-Pomares et al., 2018). We thus first analyzed the expression of EpCAM in epithelial Caco2 cells cultured on collagen-coated substrates at different culture times, *i.e.* either as 21-days monolayers or 2-days post-plating at very low density to obtain a vast majority of single cells. We observed that EpCAM was already expressed in single cells at a level comparable to monolayers (Figure 1a-b), suggesting that EpCAM may play important functions early at the single cell stage. To determine the impact of EpCAM silencing on individual cell behavior and organization, we used the stable control and EpCAM-silenced Caco2 clones that were previously established (Salomon et al., 2017) and assessed cell spreading and migration by time-lapse imaging. Right after seeding, almost all control cells were able to attach and spread while only half of EpCAM-silenced cells did so (Figure 1c). Within two hours, control cells completed spreading and spontaneously developed an elongated polarized shape before active crawling (Supplementary video 1; Figure 1d-e). In sharp contrast, mutant cells abnormally spread and failed to polarize (Supplementary video 2; Figure 1d). The isotropic organization as a circular or “fried-egg” shape, characteristic of unpolarized single cells, was stable over 2-days post seeding (Figure 1f). Quantification of the aspect ratio (Figure 1g) and the distance between the nucleus and the cell centroid (Figure 1h) further confirmed the cell polarization defect induced by the loss of EpCAM (aspect ratio of 1.76+/− 0.42 for control cells; 1.19+/− 0.13 and 1.17+/-0.11 for EpCAM shRNA#1 and #2 (mean +/− SD)). Noteworthy, a subpopulation of fried-egg shaped EpCAM-depleted cells showed a symmetry breaking event and eventually displayed a large C-shaped protrusion reminiscent of fish keratocytes migration mode (31 and 36% in Caco2 shEpCAM#1 and #2, respectively; Figure 1i, Supplementary Figure 1), as was reported in fibroblasts after ROCK1 inhibition (Cai et al., 2010). Nevertheless, these EpCAM-KD C-shaped cells do not show any directional motility, instead rotating on themselves and detaching rapidly from the substrate (Supplementary video 3). The loss of front-rear polarity in mutant cells prompted us to test their cell migratory behavior. Whereas control cells exhibit an active motile behavior, as assessed by cell displacement (Figure 1j; Supplementary video 1), the displacement of EpCAM-KD cells on the substrate was less extensive (Figure 1k-l; Supplementary video 2). Taken together, the data show that: i) EpCAM plays a role in single cells, independently of cell-cell contacts, and ii) the absence of EpCAM provokes changes early during cell morphology acquisition, generating a stable unpolarized state which impinges on the migratory behavior of epithelial cells.

**Figure 1:**
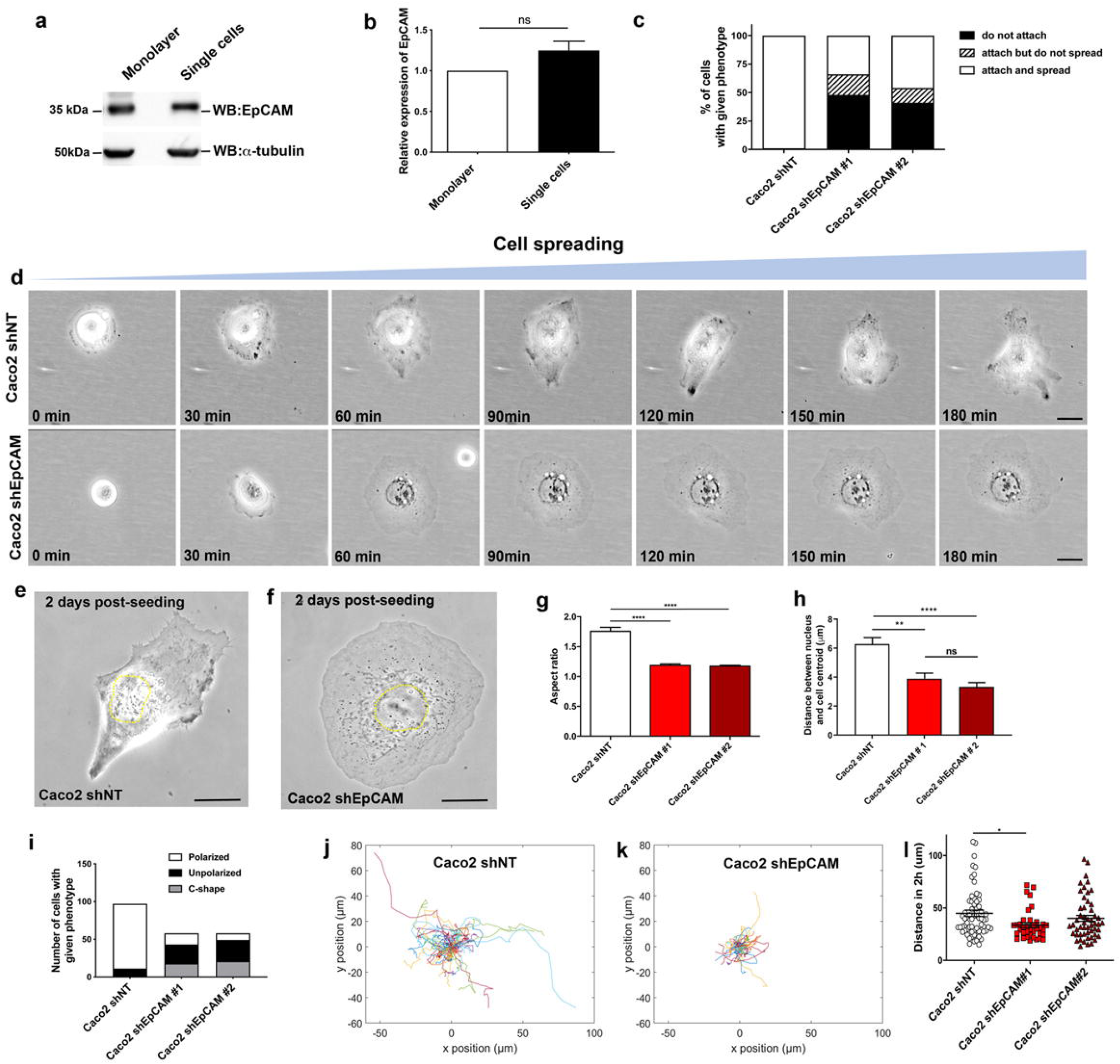
EpCAM is required for single cell front-rear polarization and migration. **(a-b)** Western blot analysis **(a)** and statistical analysis **(b)** of the EpCAM expression in Caco2 21-days monolayers or single cells spread on collagen-coated petri dish and coverslips, respectively. α-tubulin was used as a loading control. Three independent experiments were carried out. (**c**) Statistical analysis of cell adhesion within three hours post-seeding for control *(Caco2 shNT)* or EpCAM-depleted *(Caco2 shEpCAM #1 and #2)* cells. Are represented the percentage of cells that attach and spread, attach but do not spread, and cells that do not attach. N *(Caco2 shNT)* = 101 cells, N *(Caco2 shEpCAM#1)* = 121 cells, N *(Caco2 shEpCAM #2)* = 112 cells. Chi-square test computed on the number of cells with given phenotype indicates a p-value < 0.0001. Three independent experiments were carried out. (**d**) Phase contrast time-lapse of control and EpCAM-silenced cells during cell spreading. Imaging was performed for three hours right after seeding. Scale bar, 5μm. **(e-f)** Phase contrast representative images of control **(e)** or EpCAM-depleted **(f)** 2-days post-seeding Caco2 single cells. Scale bar, 5μm. **(g)** Statistical analysis of the aspect ratio in control *(Caco2 shNT)* or EpCAM-deprived *(Caco2 shEpCAM #1 and #2)* cells. Mean aspect ratio for Caco2 shNT cells = 1.756±0.06, Caco2 shEpCAM#1 = 1.188±0.0254, Caco2 shEpCAM#2 = 1.174±0.016. Data are mean +/− SEM. **(h)** Statistical analysis of the distance between the nucleus and the centroid in control *(Caco2 shNT)* or EpCAM-deprived *(Caco2 shEpCAM #1 and #2)* cells. Mean distance between the nucleus and the centroid in Caco2 shNT cells = 6.28±0.46, Caco2 shEpCAM#1 =3.911±0.38, Caco2 shEpCAM#2 = 3.321±0.32. N *(Caco2 shNT)* = 41 cells, N *(Caco2 shEpCAM#1)* = 33 cells, N *(Caco2 shEpCAM#2)* = 44 cells. Kruskal-Wallis test with Dunn’s multiple comparison test, ** adjusted *P* =0.0015, **** adjusted *P* <0.0001. Three independent experiments were carried out. Values are mean ± s.e.m. (**i**) Statistical analysis of polarity phenotypes in control *(Caco2 shNT)* or EpCAM-depleted *(Caco2 shEpCAM#1 and #2)* cells. N *(Caco2 shNT)* = 97 cells, N *(Caco2 shEpCAM#1)* = 58 cells, N *(Caco2 shEpCAM #2)* = 58 cells. Three independent experiments were carried out. (**j-k**) Color map of cell tracks in control *(Caco2 shNT)* **(j**) and EpCAM-depleted *(Caco2 shEpCAM)* **(k)** cells. **(l)** Statistical analysis of the distance traveled by control and EpCAM-silenced cells within 2 hours. N *(Caco2 shNT)* = 67 cells, N *(Caco2 shEpCAM#1)* = 37 cells, N *(Caco2 shEpCAM #2)* = 53 cells. ANOVA test, * *P*-value=0,03. Values are mean ± s.e.m. Three independent experiments were carried out.

### Block in stress fiber maturation in the absence of EpCAM

Many studies reported that acquisition of single cell polarity is achieved through changes in the ordering of actin cytoskeleton and adhesive structures (Geiger et al., 2009; Prager-Khoutorsky et al., 2011; Ladoux et al., 2016; Gupta et al., 2019). In line with this idea, monitoring the actin dynamics during spreading and polarity acquisition revealed that in control epithelial cells, actin cables dynamically reorganize while the cells spread and acquire a front-rear axis. First, circumferential arcs form, at the boundary between lamellipodium and lamella, coupled with the appearance of radial fibers within 30 min, which then give rise to stress fibers within 2 hours, as previously described in fibroblasts (Shemesh et al., 2009; Burnette et al., 2011, 2014) (Figure 2a, Supplementary video 4). However, although EpCAM-KD cells generated circumferential arcs and radial fibers, they kept this actin cable organization in a seemingly frozen state during the 2-hour course of the experiment (Figure 2a, Supplementary video 5), suggesting that the process of stress fiber formation may be impaired in mutant cells. Accordingly, we first analyzed the actin network architecture 2-days post seeding together with the distribution of the focal adhesion marker paxillin. While control cells displayed a majority of stress fibers (Figure 2b-c), EpCAM-silenced cells exhibited very few, instead containing a dense central network of circumferential arcs and longer radial fibers compared to control cells (Figure 2b-d). We then analyzed the co-distribution of α4-actinin and myosin-IIA. Stress fibers in control cells are cross-linked by a periodic distribution of α4-actinin that alternates with myosin-IIA (Supplementary Figure 2a). By contrast, in EpCAM-KD cells, α4-actinin accumulated on radial fibers devoid of myosin-IIA, which is enriched along circumferential arcs (Supplementary Figure 2a), in agreement with the canonical radial fibers and circumferential arcs as described in fibroblasts and osteosarcoma cell line (Cai et al., 2010; Burridge and Wittchen, 2013). Coincidently, as the major fibers in EpCAM-KD cells are radial, the connected FAs are radially oriented and mainly located in a 5μm belt at the cell periphery (Figure 2e). However, FA’s length was only slightly increased in mutant cells (Supplementary Figure 2c-d), and β1-integrin, the tension-sensitive proteins talin and vinculin, and zyxin were still located at radially-oriented FAs in the absence of EpCAM (Supplementary Figure 2b), showing that EpCAM’s loss barely impacts FA’s composition or morphology *per se* but rather their location. The specificity of these abnormalities was tested with rescue experiments by transfecting an EpCAM-GFP shRNA-resistant construct in EpCAM-depleted cells. Correct actin and FA localization was recovered after EpCAM rescue (Supplementary Figure 3a). These results show that EpCAM depletion perturbs actin organization and subsequently FA location in single epithelial cells, and suggest that stress fiber formation may be impaired in the absence of EpCAM.

**Figure 2:**
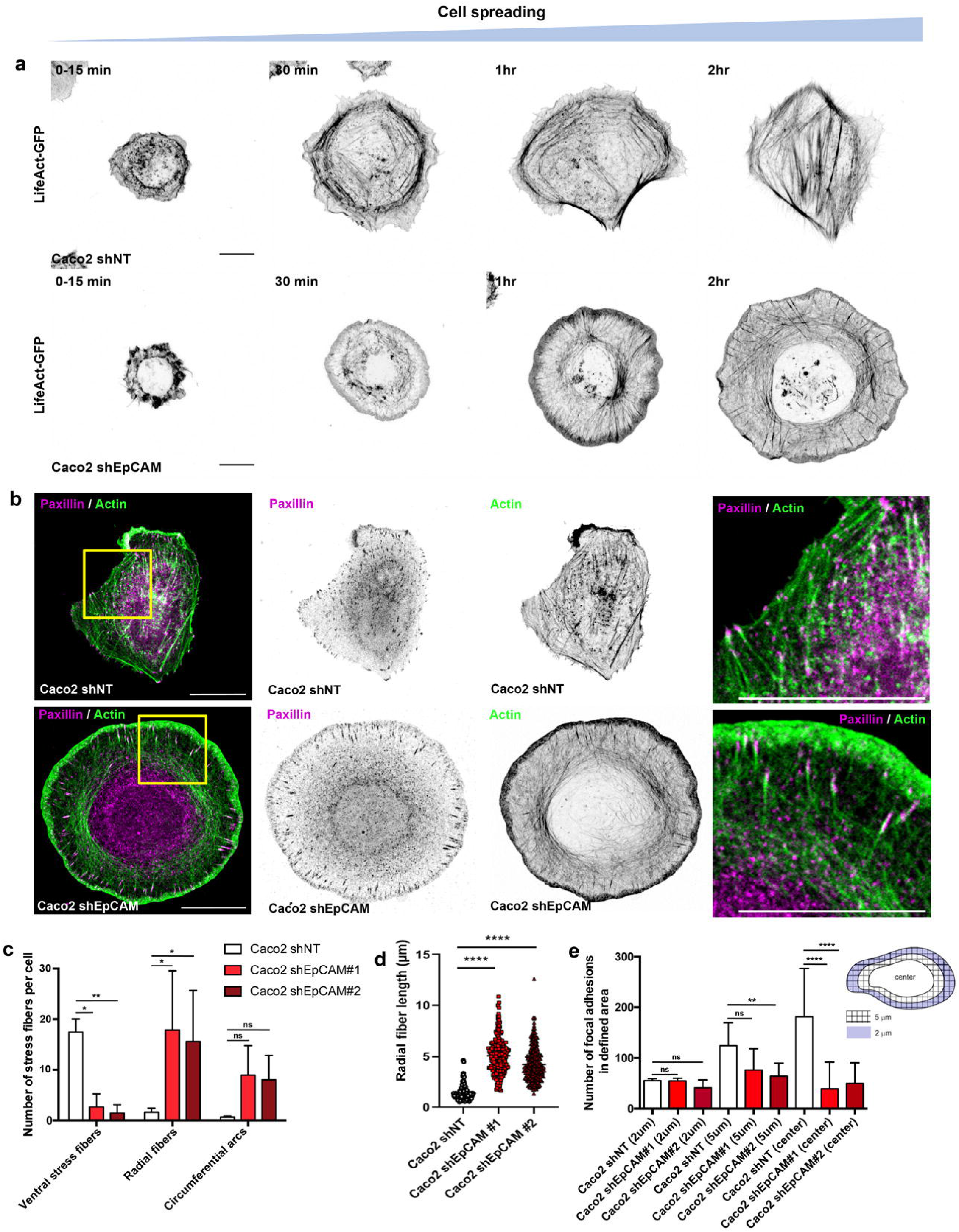
EpCAM participates in the maturation of ventral stress fibers in Caco2 cells. **(a)** Timelapse images of actin cable rearrangement during cell spreading and polarity acquisition in LifeAct-GFP transfected control and EpCAM-KD cells. Scale bar, 5μm. **(b)** Confocal analysis of actin *(green)* and paxillin *(magenta)* distributions in control *(Caco2 shNT)* and EpCAM-silenced *(Caco2 shEpCAM)* cells. Areas boxed in yellow are presented on the right. Accumulated z-stack are presented. Scale bar, 5μm. **(c)** Statistical analysis of the number of ventral stress fibers, radial fibers and circular arcs in control and EpCAM-depleted cells. N *(Caco2 shNT)* = 30 cells, N *(Caco2 shEpCAM#1)* = 30 cells, N *(Caco2 shEpCAM #2)* = 30 cells. Sidak’s multiple comparisons test, \**P* <0.02, \*\**P* <0.0011. **(d)** Statistical analysis of the length of radial fibers in control and EpCAM-silenced cells. N *(Caco2 shNT)* = 36 cells, N *(Caco2 shEpCAM#1)* = 35 cells, N *(Caco2 shEpCAM #2)* = 43 cells, n>160 dorsal fibers for each condition. Kruzkal-Wallis test and Dunn’s multiple comparison test, \*\*\*\**P*<0.0001. Values are mean ± s.e.m. **(e)** Statistical analysis of FA density in the delimited 2μm, 5μm from the cell periphery or center area. Mean number of FA in the 2μm area for Caco2 shNT cells = 55.03±4.12, Caco2 shEpCAM#1 = 54.50±5.26, Caco2 shEpCAM#2 = 40.93±2.86, in the 5μm area for Caco2 shNT cells = 124.4±8.13, Caco2 shEpCAM#1 = 76.40±7.68, Caco2 shEpCAM#2 = 63.97±4.71, in the center area for Caco2 shNT cells = 181.5±17.07, Caco2 shEpCAM#1 = 38.87±9.67, Caco2 shEpCAM#2 = 49.73±7.47. N *(Caco2 shNT)* = 31 cells, N *(Caco2 shEpCAM#1)* = 30 cells, N *(Caco2 shEpCAM #2)* = 30 cells. Kruzkal-Wallis test and Dunn multiple comparison test, \*\**P* <0.01, ****P<0.0001. For each experiment, three independent experiments were carried out.

### Loss of EpCAM modifies cell mechanical properties

Given the stress fiber subtype differences in tension-bearing and force generation properties (Lee et al., 2018), we reasoned that the distinct actin organization could account for a modification of the cell mechanical properties and would explain the defective migratory behavior of EpCAM-KD cells. Atomic force microscopy experiments showed that similar rigidity was detected in the central region containing the nucleus, *i.e.* in cell heights range comprised between 2 and 5μm, in control and mutant cells (Figure 3a-b). However, as expected, different rigidities were measured in the cell protrusion, considered as the area where the cell height is below 2μm (two-fold increase in Young’s modulus for EpCAM-KD cells in the cell area under 2μm in height) (Figure 3a,c), demonstrating that EpCAM depletion resulted in a higher cortical stiffness of the protrusion. This difference is most likely due to the higher density of contractile circumferential fibers in the EpCAM-KD cells resolved by the AFM nanoindentation (Figure 3a). This finding suggests that EpCAM may play a role in regulating intracellular stiffness through its action on the actin cytoskeleton, its depletion leading to stiffer cells, less deformable and thus less polarized, as previously suggested for rigidity sensing mechanism (Trichet et al., 2012).

**Figure 3:**
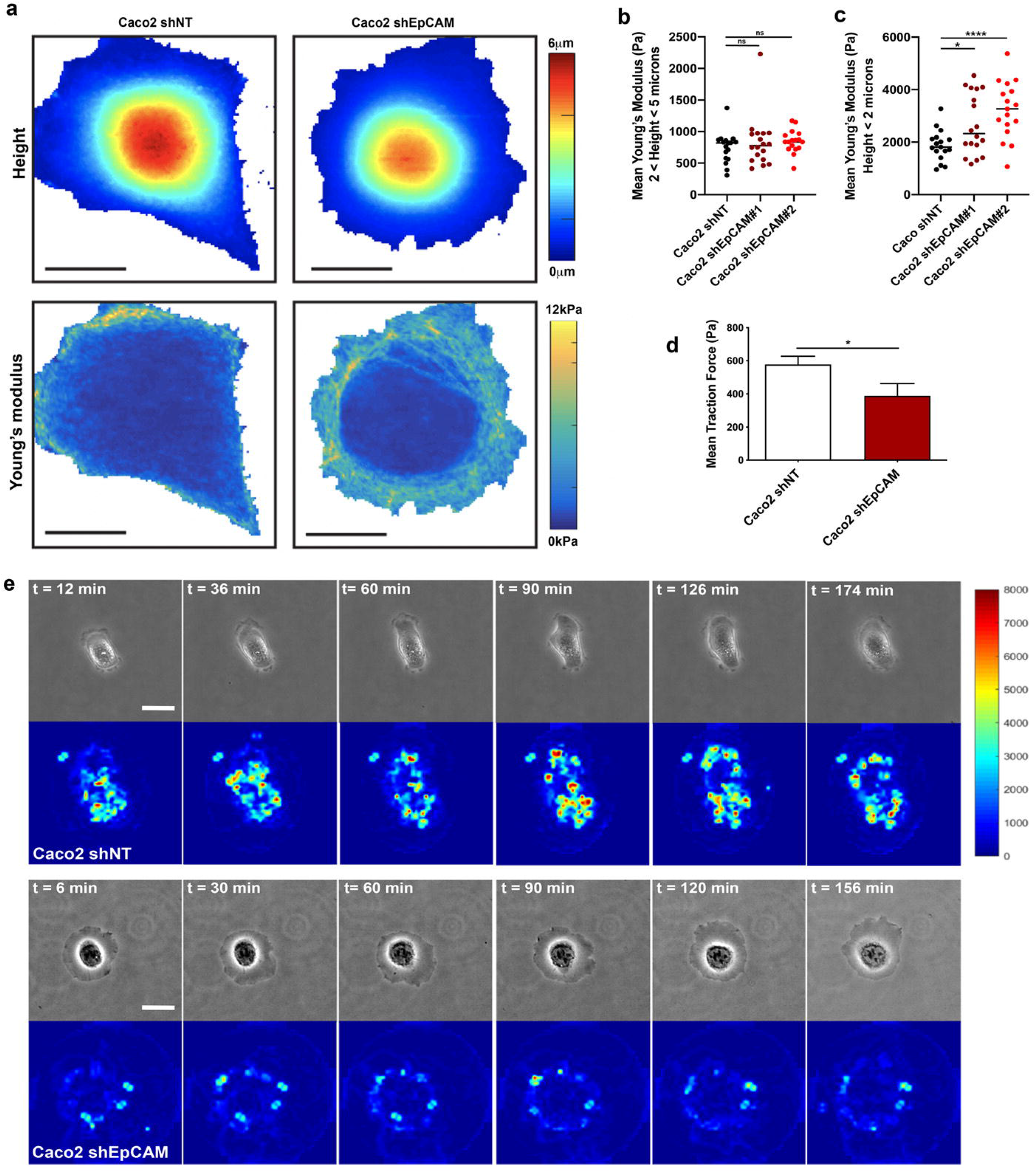
Loss of EpCAM provokes cell mechanical changes. (**a**) Representative maps showing the cell topography *(top)* and local Young’s modulus *(bottom)* for a representative control cell *(left)* and shEpCAM cell (*right*). Scale bar, 20μm. (**b-c**) Average Young’s modulus in the region of height in the range of 2 to 5 μm (**b**) or in the region of height < 2 μm (**c**) for each cell. (N *(Caco2 shN*T) = 13 cells, N *(Caco2 shEpCAM#1)* = 14 cells, and N *(Caco2 shEpCAM #2)* = 13 cells. Unpaired t-test, * *P* =0.0149, **** *P* <0.0001. Three independent experiments were carried out. **(d)** Statistical analysis of the mean traction forces measured in control *(Caco2 shNT)* and EpCAM-silenced *(Caco2 #2)* cells. N *(Caco2 shNT)* = 40 cells, N *(Caco2 shEpCAM #2)* = 13 cells. Student test, \**P* = 0.0236. Three independent experiments were carried out. (**e**) Representative phase contrast and color map images of traction forces exerted by control *(Caco2 shNT)* and EpCAM-deprived *(Caco2 shEpCAM)* cells. Scale bar, 5μm.

Furthermore, traction force microscopy (TFM) experiments (Trepat et al., 2009) allowed measurement of the impact of actin remodeling in mutant cells on their ability to generate traction forces on the substrate. The data revealed that EpCAM-KD cells exerted lower traction forces on the substrate than the control cells (mean of 388.1 ± 75.0 Pa and 577.4 ± 49.9 Pa, respectively) (Figure 3d). Additionally, time-lapse analysis of TFM data showed that control cells generate traction in a dynamic manner, probing the substrate, while EpCAM-KD lower forces are maintained throughout the experiment (Figure 3e). Thus, changes in the organization of the actin cytoskeleton between normal and mutant cells impact the level of traction forces as well as its distribution. Consequently, the absence of contractile stress fibers at FA sites in EpCAM-KD leads to lower forces as previously observed in similar cases (Ladoux et al., 2016). Altogether, the data demonstrate that EpCAM deprivation modifies the mechanical properties of single epithelial cells.

### EpCAM expression potentiates stress fiber formation

EpCAM is expressed in several simple epithelia, and we wondered whether its impact on stress fiber organization could be observed in cell assemblies. Similar mis-arrangement of FAs and actin cables was observed in EpCAM-KD cell islands (Supplementary Figure 3b), showing that the impact of EpCAM on cell-substrate adhesion and actin organization also takes place when cell-cell contacts are formed. To investigate if this effect of EpCAM expression could be generalized, we probed the stress fiber organization of other EpCAM-expressing or non-expressing cell types, reasoning that they would behave as our control, or EpCAM-deprived clones, respectively. We used the renal epithelial MDCK cells, which display an EpCAM expression level comparable to control Caco2 cells, and osteosarcoma U2OS cells and endometrial epithelial HeLa cells, showing similar lack of EpCAM expression as EpCAM-depleted Caco2 clones (Figure 4a, b). Whereas control MDCK cells exhibited tangential stress fibers, their formation was impaired upon EpCAM-siRNA silencing (Figure 4c, f; Supplementary Figure 4). Conversely, while U2OS and HeLa cells display a vast majority of radial and transverse arcs, as previously reported (Burridge and Wittchen, 2013), the introduction of an EpCAM-GFP construct induced a remodeling towards stress fibers, demonstrating that ectopic EpCAM expression is sufficient to drive actin fiber rearrangement in EpCAM non-expressing cells (Figure 4d-e, g-h). We conclude that EpCAM participates to a cell-autonomous general regulatory mechanism for cell polarity, by potentiating stress fiber maturation.

**Figure 4:**
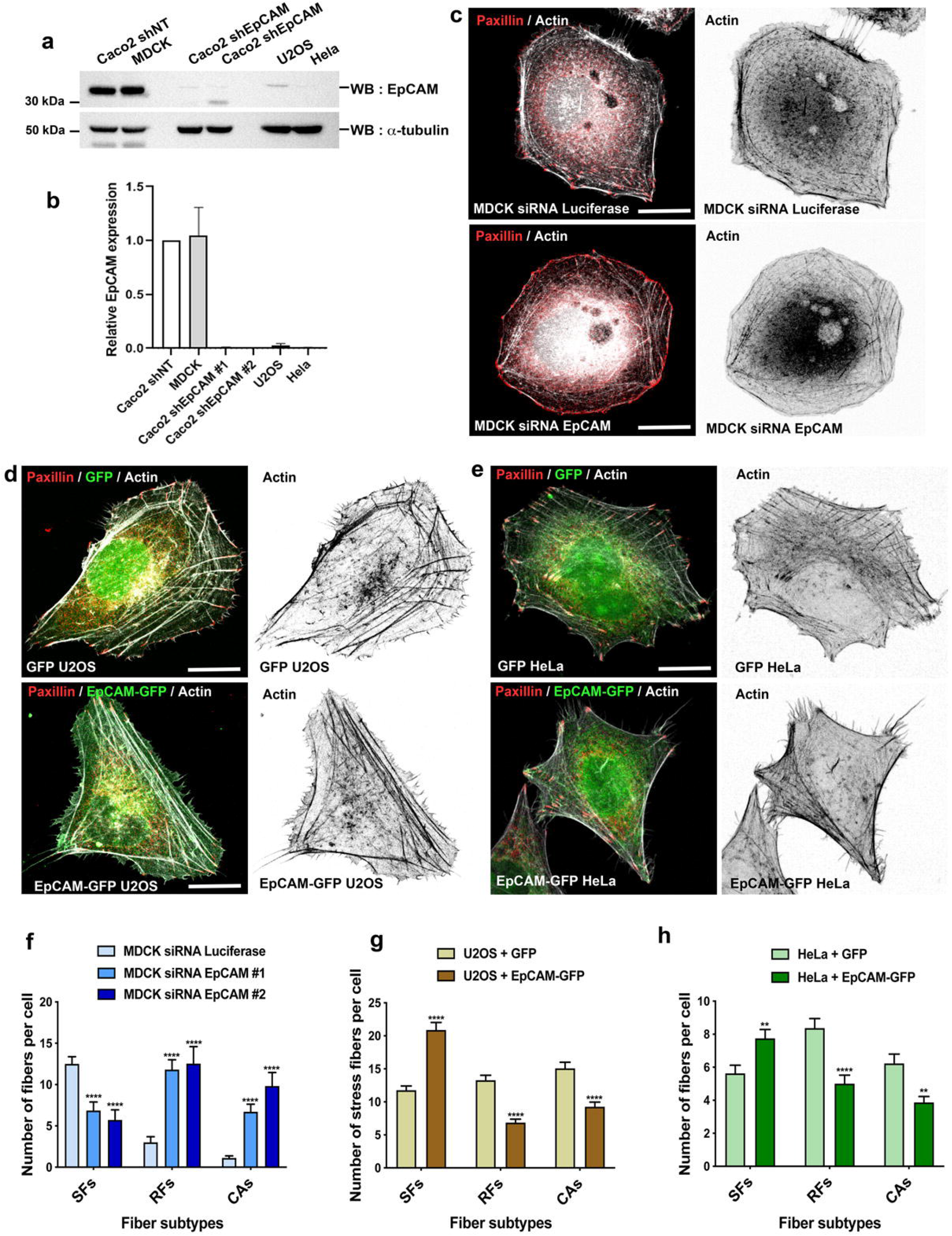
EpCAM expression triggers the formation of stress fibers. **(a)** Western blot analysis of EpCAM expression in control Caco2 *(Caco2 shNT),* MDCK, EpCAM-silenced *(Caco2 shEpCAM#1 and #2),* U2OS and HeLa cells. α-tubulin was used as a loading control. **(b**) Quantification of EpCAM expression in control Caco2 *(Caco2 shNT),* MDCK, EpCAM-silenced *(Caco2 shEpCAM#1 and #2),* U2OS and HeLa cells. Three independent experiments were carried out. **(c-e)** Confocal analysis of actin and paxillin in control and EpCAM siRNA-treated MDCK cells **(c)**, in GFP- and EpCAM-GFP-transfected U2OS cells **(d)**, and in GFP- and EpCAM-GFP-transfected HeLa cells **(e)**. Accumulated z-stack are presented. Scale bar, 5μm. (**f**) Statistical analysis of the number of stress fibers (SFs), radial fibers (RFs) and circumferential arcs (CAs) in control (siRNA Luciferase) or EpCAM-silenced (siRNA EpCAM) MDCK cells. N *(MDCK siRNA Luciferase)* = 33 cells, N *(MDCK siRNAEpCAM#1)* = 21 cells, N *(MDCK siRNAEpCAM#2)* = 11 cells. Multiple t-test, **** *P*<0.0001. (**g**) Statistical analysis of the number of stress fibers, radial fibers and circular arcs in U2OS cells transfected with either GFP (*GFP U2OS)* or EpCAM-GFP *(EpCAM-GFP U2OS).* N (*GFP U2OS)* = 50 cells, N *(EpCAM-GFP U2OS)* = 58 cells. Multiple t-test, **** *P*<0.0001. (**h**) Statistical analysis of the number of stress fibers, radial fibers and circular arcs in HeLa cells transfected with either GFP *(GFP HeLa)* or EpCAM-GFP *(EpCAM-GFP HeLa).* N (*GFP HeLa)* = 36 cells, N *(EpCAM-GFP HeLa)* = 44 cells. Multiple t-test, ** *P*<0.01, **** *P*<0.0001. For each experiment, three independent experiments were carried out.

### Aberrant contractile activity is at the origin of stress fiber formation and cell polarity failure

Several mechanisms were put forward to explain stress fiber maturation. The striking phenotype developed by EpCAM-KD cells prompted us to investigate the participation of α4-actinin and myosin-IIA. α4-actinin first appeared as a candidate of choice, since it has been reported in literature as an EpCAM binding partner (Balzar et al., 1998). However, in our hands, no interaction between EpCAM and α4-actinin was detected using co-immunoprecipitation (Supplementary Figure 5a). In addition, α4-actinin silencing by siRNA was not able to restore the presence of stress fibers in the EpCAM-KD cells (Supplementary Figure 5b-c). Previous studies described that the intensity and distribution of cell contractility were modulated by EpCAM in epithelial assemblies (Maghzal et al., 2013; Salomon et al., 2017). We therefore focused on cell contractility mechanisms to determine whether they are involved in the development of the EpCAM-KD phenotype in isolated cells. As shown in Supplementary Figure 2a, myosin-IIA remains associated with circumferential arcs in mutant cells. To assess the contractile ability of myosin-IIA, we performed an immunostaining against the phosphorylated form of the myosin regulatory light chain (P-MLC2). In absence of EpCAM, the P-MLC2 signal intensified, as described previously in Caco2 cell clusters (Magzhal et al., 2013). Moreover, the P-MLC2 signal was concentrated along circumferential arcs in comparison to control cells (Figure 5a-b). This result was confirmed by Western blot, where P-MLC2 levels increased relative to the total amount of MLC2 in comparison with control cells (Figure 5c-d). These data demonstrate that actomyosin activity is increased in single EpCAM-silenced cells but restricted to circumferential actin arcs, creating a uniform hypercontractile ring at the cell cortex.

**Figure 5:**
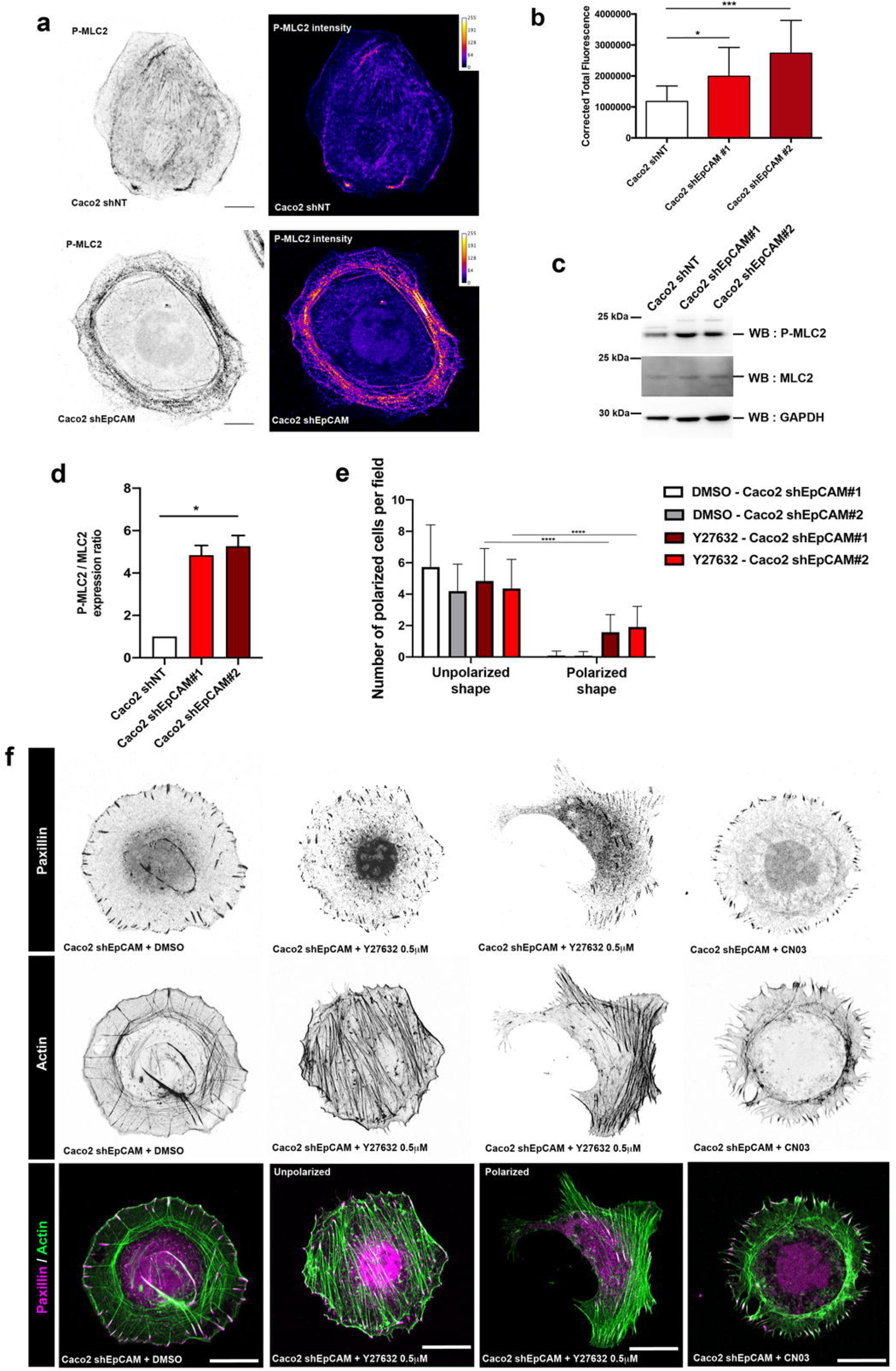
Defective cell contractility activity and distribution is responsible of the development of the EpCAM-KD phenotype. **(a)** Confocal analysis of the distribution of P-MLC2 in control and EpCAM-silenced Caco2 cells. Accumulated z-stack are presented. Color coded images for P-MLC2 intensity signal are presented on the right panel. Scale bar, 5μm. **(b)** Statistical analysis of the corrected total fluorescence for P-MLC2 in control *(Caco2 shNT)* and EpCAM-silenced *(Caco2 shEpCAM #1 and #2)* cells. *(Caco2 shNT)* = 10 cells, N *(Caco2 shEpCAM#1)* = 10 cells, N *(Caco2 shEpCAM #2)* = 10 cells. One-way ANOVA with unpaired t-test, * *P*-value < 0,0252, *** *P*-value <0,0005. **(c)** Western blot analysis of MLC and P-MLC amounts in control *(Caco2 shNT)* and EpCAM-silenced *(Caco2 shEpCAM #1 and #2)* Caco2 cells. GAPDH was used as a loading control. **(d)** Statistical analysis of P-MLC2 amount relative to MLC2 in control and EpCAM-depleted cells. Kruskal-Wallis test, * *P*-value = 0.04. Three independent experiments were carried out. **(e)** Statistical analysis of the number of unpolarized- and polarized-shaped Caco2 shEpCAM#1 and #2 cells after DMSO or Y-27632 0.5 μM treatment. N (DMSO Caco2shEpCAM#1) = 192 cells, N (DMSO Caco2shEpCAM#1) = 111 cells, N (Y-27632 Caco2shEpCAM#1) = 199 cells, N (Y-27632 Caco2shEpCAM#2) = 207 cells. Paired t-test, **** *P*<0.0001. Three independent experiments were carried out. (**f**) Confocal analysis of paxillin *(magenta)* and actin (*green*) in EpCAM-depleted cells upon DMSO, blebbistatin 2μM, Y-27632 0.5 μM, CN03 1 μg/ml or SMIFH2 2μM treatment for 1 hour. Accumulated z-stack are presented. Scale bar, 5μm.

To test whether the local hyperactivity of the actomyosin network was at the origin of stress fiber and FA abnormalities and to identify the involved signaling pathway, we submitted EpCAM-KD cells to diverse drug treatments affecting cell contractility and actin polymerization. Reducing myosin-II ATPase activity using incubation with a classical 10μM blebbistatin dose led to total disappearance of radial fibers and circular arcs, as previously described (Supplementary Figure 6a) (Burnette et al., 2014; Tee et al., 2015). However, treatment with 2 μM blebbistatin resulted in a decreased number of circumferential arcs, the swirling of actin cables with the development of few stress fiber, as well as more numerous and more centrally distributed FAs (Supplementary Figure 6a). Similarly, treatment with a classical 10 μM Y27632 dose to reduce ROCK activity totally abolished the formation of radial fibers and circumferential arcs (Supplementary Figure 6a), but a low dose (Y27632, 0.5 μM) led to the disappearance of radial stress fibers, the formation of linear stress fibers and the FA redistribution within the cell body (Figure 5f). In addition, almost one third of the EpCAM-KD treated cells recovered a polarized shape after Y-27632 low dose treatment (Figure 5e-f). We concluded that mild adjustments of the level of circumferential arc contractility is able to trigger partial or full recovery of the stress fiber maturation as well as recovery of polarized shape using blebbistatin and Y-27632 treatment, respectively. Interestingly, these data pointed towards RhoA signaling. To further assess the upstream RhoA involvement, we directly evaluated the level of endogeneous active RhoA by FRET experiment in control or EpCAM-silenced cells. We used the FRET probe developed by Matsuda and colleagues (Yoshizaki et al., 2003), consisting of YFP, Rhotekin-RBD and CFP (Supplementary Figure 6b). EpCAM silencing led to a significant decrease of FRET ratio, testifying of an increase of RhoA activity in the absence of EpCAM (Supplementary Figure 6c-d). Accordingly, the over-activation of RhoA by treating mutant cells with RhoA activator (CN03) dramatically worsened the EpCAM-KD phenotype (Figure 5f). Moreover, EpCAM-KD cells display longer radial stress fibers (Figure 2d). This data prompted us to test the participation of formins since they are involved in radial fiber polymerization and are also well-known effectors of RhoA (Kühn and Geyer, 2014). Along this line, we evaluated their activity in EpCAM-depleted cells using the SMIFH2 inhibitor. SMIFH2 low-dose treatment reduced the size of the radial fibers, as well as the cell protrusion depth (Supplementary Figure 6e) but failed to restore stress fiber development and cell shape remodeling. It is worth mentioning that a treatment with the Rac1 inhibitor NSC23766 strongly reduced the size of the lamellipodium but failed to rescue correct actin cable and adhesive structure arrangements (Supplementary Figure 6e). Moreover, Arp2/3 or MLCK inhibition, through CK666 or ML-7 treatment, respectively, had no effect on actin cytoskeleton arrangement in EpCAM-silenced cells (Supplementary Figure 6e). Together, the findings testified of a main participation of RhoA signaling towards cell contractility regulation rather than actin polymerization in the EpCAM-dependent mechanism. We concluded that local actomyosin hyperactivity is at the core of the defects on stress fiber development and polarity acquisition induced by the silencing of EpCAM, and we hypothesized that EpCAM may act on upstream actomyosin apparatus activity, probably at the level of RhoA signaling.

### Cortical active RhoA zone is remodeled during epithelial cell spreading and required EpCAM

To determine the impact of EpCAM on Rho signalling, we transfected Caco2 cells with either wt RhoA (GFP-Rho), a constitutively active mutant (GFP-RhoG14V) or a dominant negative mutant of RhoA (GFP-RhoT19N). In control cells, expression of mutant forms of Rho destabilized actin network and FA organization, as expected (Supplementary Figure 7). Whereas the expression of GFP-RhoG14V led to an increase of FA number and stress fiber formation, the expression of GFP-RhoT19N provoked a reduction of FA number, their concentration at the cell periphery and a decrease in stress fiber content (Supplementary Figure 7). Thus, the expression of both mutant forms of Rho only partially recapitulate parts of the EpCAM-KD phenotype in control cells. In addition, the introduction of GFP-RhoG14V worsened the phenotype of EpCAM-silenced cells, resembling CN03 treated cells (Figure 5f). After GFP-RhoT19N transfection, no obvious change was observed for actin arrangement and FAs in EpCAM-KD cells (Supplementary Figure 7). Together these results led us to conclude that RhoA activity may contribute to the development of defects under EpCAM silencing. Since gross RhoA modulation through the use of constitutively active or inactive mutants seems insufficient to explain the EpCAM-KD phenotype, we hypothesized that a spatial and/or a temporal factor might be missing in this analysis.

We thus decided to assess the subcellular localization of the GTP-loaded form of RhoA by taking advantage of the fluorescent location biosensor derived from the C-terminus of anillin, AHPH (Tse et al., 2012; Priya et al., 2015). mCherry-tagged AHPH partially overlaps with total Rho-GFP (Supplementary Figure 8a) and significantly co-distributes with location biosensors of the RhoA effectors ROCK1 and mDia (ROCK1-GBD-GFP, based on the GTP-RhoA binding domain of ROCK1, and mDia-GBD-GFP, based on the GTP-RhoA binding domain of mDia) (Supplementary Figure 8b-c) (Budnar et al., 2019). Moreover, the expression of an AHPH mutated form in the RBD domain, unable to bind GTP-RhoA (AHPH^A740D^-GFP), generates a different intracellular pattern than the wt AHPH form (Supplementary Figure 8d) (Priya et al., 2015; Vassilev et al., 2017). These data testify of the specificity of AHPH-tagged for RhoA-GTP signal and confirm that it can be faithfully used to probe RhoA-GTP dynamics. We first scrutinized the spatial distribution of RhoA together with its GTP-loaded form (Rho-GFP and AHPH-mCherry, respectively; Figure 6a). Whereas control cells display partial colocalization of RhoA-GFP and AHPH-mCherry in intracellular structures, as previously reported in endothelial and neuronal cells (Bisi et al., 2013; Braun et al., 2015; Vassilev et al., 2017), EpCAM-KD cells displayed an accumulation of both RhoA-GFP and its active form in large tubular compartments within the lamella (Figure 6a-b, c), where their colocalization drastically increases up to 70% (Figure 6b). These results suggest that a slow-down in the GTPase cycling might occur, keeping the GTP-RhoA form longer-lived in absence of EpCAM. This hypothesis would be supported by a block of RhoA dynamics in tubular compartments. We followed the reporter’s behavior, and the large spread of GTP-RhoA displacement patterns revealed a complex intracellular dynamics in the cell protrusion of control cells (Figure 6d-e; Supplementary video 6). In the absence of EpCAM however, GTP-RhoA movement was extremely impaired with a speed decrease and reduced displacement (Figure 6d-h; Supplementary video 7). Transfection the EpCAMr-GFP shRNA-resistant construct in EpCAM-silenced cells restored a correct distribution for AHPH (Supplementary Figure 9a). These data showed that EpCAM is required for the correct dynamics of RhoA-GTP in the cell protrusion.

**Figure 6:**
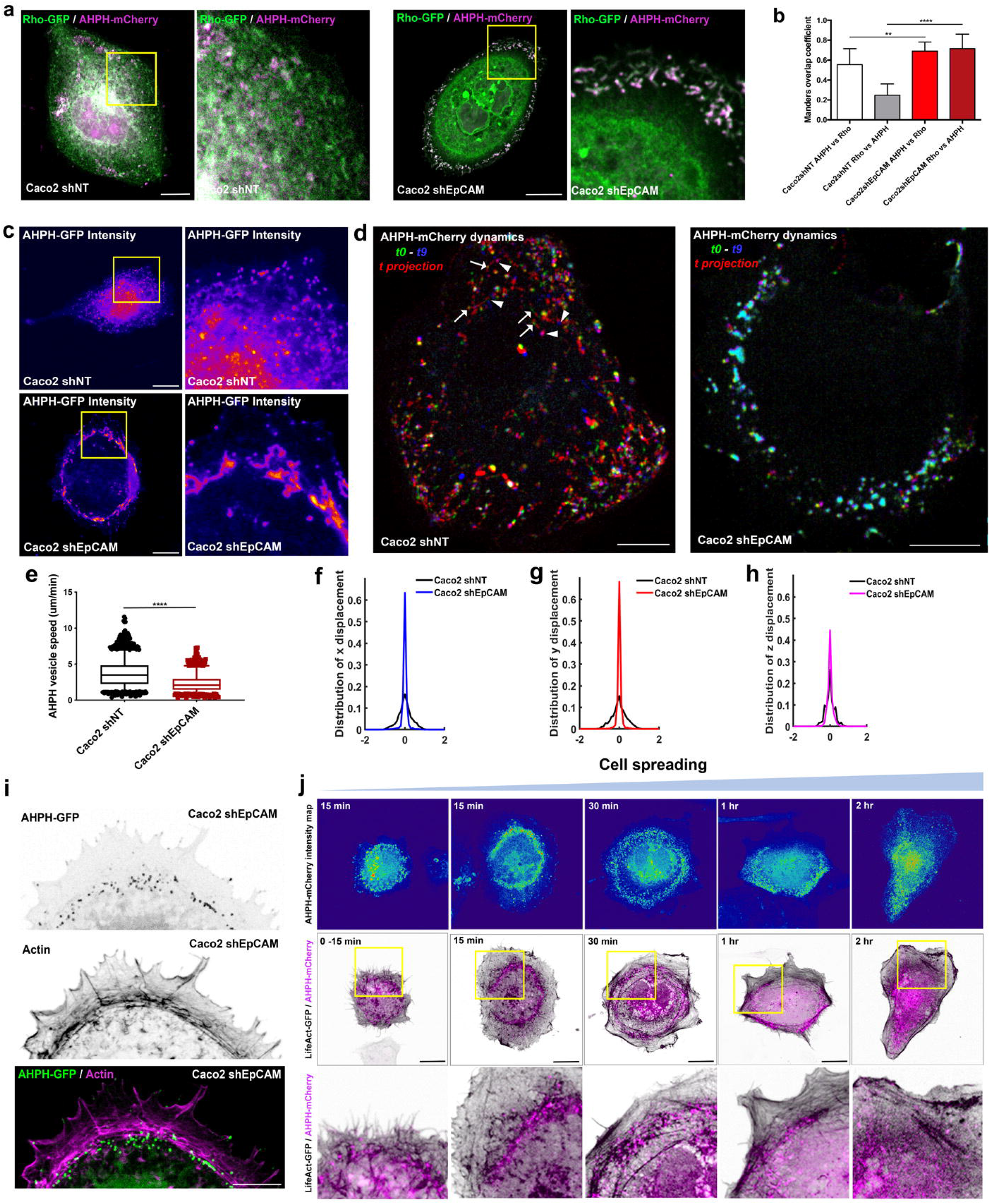
Active RhoA distribution is impaired in the absence of EpCAM. **(a)** Confocal analysis of the distribution of RhoA-GFP (*green*) together with AHPH-mCherry *(magenta)* in control and EpCAM-silenced cells. Areas boxed in yellow are presented on the right. Accumulated z-stack are presented. Scale bar, 5μm. (**b**) Quantification of the proportion of the AHPH-mCherry probe overlapping with total RhoA. Manders overlap coefficient for AHPH-mCherry versus RhoA-GFP in Caco2 shNT cells = 0.5558±0.04, and in Caco2 shEpCAM cells = 0.6913±0.02. Manders overlap coefficient for RhoA-GFP versus AHPH-mCherry in Caco2 shNT cells = 0.2485±0.03, and in Caco2 shEpCAM cells = 0.7165±0.03. N *(Caco2 shNT)* = 5 cells, N *(Caco2 shEpCAM #1)* = 5 cells. T-test, \*\**P* = 0.0049; \*\*\*\**P* <0.0001. Values are mean ± s.e.m.. **(c)** AHPH-GFP intensity maps were generated in control and EpCAM-depleted cells with LUT table Physics from ImageJ. Areas boxed in yellow are presented on the right. Accumulated z-stack are presented. Scale bar, 5μm. (**d**) Color-coded t-projection of 10 frame time-lapse series of AHPH-mCherry in control and EpCAM-KD cells. The first image (*t0*) is false-colored green, the last image (*t9*) is false-colored in blue, and the intervening time points *(t2-8)* are submitted to t-projection and shown in red (*t projection*). Areas boxed in yellow are presented on the right. Arrows point at the position of some AHPH compartment at the beginning of the time-lapse series, whereas the arrowheads point to the position of the corresponding AHPH compartment at the end of the time-lapse series. Scale bars, 10μm. (**e**) Statistical analysis of the speed of AHPH-mCherry compartments in control and EpCAM-KD cells. BoxPlot line is median. Whiskers are 5-95 percentile. n *(Caco2 shNT)* = 1827 AHPH-positive vesicles, n *(Caco2 shEpCAM)* = 1234 AHPH-positive compartments. Mann-Whitney test, **** *P*-value < 0.0001. Three independent experiments were carried out. (**f-h**) Analysis of the displacement of the AHPH-mCherry compartments in the *x* (**g**), *y* (**h**) and *z* (**i**) direction in control or EpCAM-KD cells. (**i**) Confocal analysis of the distribution of AHPH-GFP (*green*) and actin cables *(magenta)* in the protrusion of EpCAM-silenced cells. Accumulated z-stack are presented. Scale bar, 5μm. **(j)** Confocal analysis of the distribution of AHPH-mCherry *(magenta)* and actin *(black)* in control Caco2 cells during cell polarization and maturation of stress fibers from 0 to 2 hours. Areas boxed in yellow are presented on the bottom panel. AHPH-mCherry intensity maps are presented in the upper panel. Accumulated z-stack are presented. Scale bar, 5μm.

Furthermore, in EpCAM-KD cells, AHPH-positive compartments were subsequently enriched in the proximity of the circumferential actin cables (Figure 6i), prompting us to envisage a positive correlation between GTP-RhoA dynamics and actin cable remodeling in control cells. Although typical transitions in cytoskeletal rearrangement during cell spreading and polarity acquisition are described, a global spatial and temporal analysis of the contractility signaling in this context is still missing. By establishing the spatiotemporal dynamics of AHPH-mCherry, we determined that active RhoA displays drastic distribution changes during cell shape remodeling (Figure 6j). Within the first 15 minutes post-seeding, whereas no clear actin cable can be distinguished yet, GTP-RhoA-positive compartments are perinuclearly located in still round-shaped cells. While spreading intensifies with the clear expansion of protrusions in circular shaped cells, GTP-RhoA concentrates in a central ring and co-distributes with an intense actin worm meshwork. The time of 30-minutes post-seeding is characterized by the appearance of circumferential arcs at the level of the active RhoA central ring. Starting from 1 hour after initiation of spreading, GTP-RhoA distribution is remodeled, causing the disappearance of the central ring zone and exhibiting a scattered and homogeneous patterning throughout the cytoplasm, concomitantly with the acquisition of stress fibers and a polarized cell shape. In conclusion, a correlation indeed exists between GTP-RhoA dynamics and actomyosin rearrangement. Altogether, these data led us to conclude that the reorganization of the cortical active RhoA zone allows the remodeling of actin fibers and the polarized cell reshaping, a pivotal step which is blocked in absence of EpCAM.

### EpCAM ensures endosomal turnover of active RhoA in the cell protrusion

Collectively, the data raised a basic question: how does EpCAM ensure proper cell spreading and actin fiber organization? In the light of afore described data, we hypothesized that EpCAM may directly act on the remodeling of the active RhoA zone during spreading. Analyzing EpCAM’s distribution together with the GTP-RhoA location biosensor, we found that a subpopulation of endogenous EpCAM- or EpCAM-GFP-positive intracellular compartments co-distributes with the RhoA reporter (Figure 7a-d, e; Supplementary Figure 9). This colocalization takes place in unpolarized cells and continues when cells acquire front-rear polarization (Figure 7a-b and 7c-d, respectively), testifying of a tight interplay between EpCAM and GTP-RhoA during cell spreading. To determine where this cooperation occurs and to go further in the comprehensive analysis of active RhoA dynamics, we screened candidate compartments and we notably used fluorescently tagged Rab GTPases as a proxy for organelle identity. For instance, EpCAM and GTP-RhoA were only barely detected in Rab5-, EEA1-, Rab7- or Rab4-positive organelles (not shown). In addition, the canonical recycling marker Rab11 only displayed 15% of colocalization with AHPH-positive compartments in Caco2 cells (Supplementary Figure 9b-c), in contrast with neural crest cells (Vassilev et al., 2017). But interestingly, we identified a preferential distribution of GTP-RhoA and EpCAM in a specific sub-fraction of endosomes controlled by Rab35 and C-terminal Eps15 homology domain-1 (EHD1). Both Rab35 and EHD1 function in fast-endocytic recycling at the level of cortical endosomes, in the early and late steps respectively (Caplan et al., 2002; Kouranti et al., 2006; Cai et al., 2013; Kobayashi and Fukuda, 2013; Kobayashi et al., 2014; Klinkert and Echard, 2016). In fact, RhoA-GTP and EpCAM co-distribute at 90% in Rab35-positive and at 60% in EHD1-positive compartments (Figure 7f, h and 7g, i, respectively). These results showed that GTP-RhoA and EpCAM co-evolve at the cell cortex and prompted us to suggest that EpCAM dictates the endosomal turnover of RhoA-GTP.

**Figure 7:**
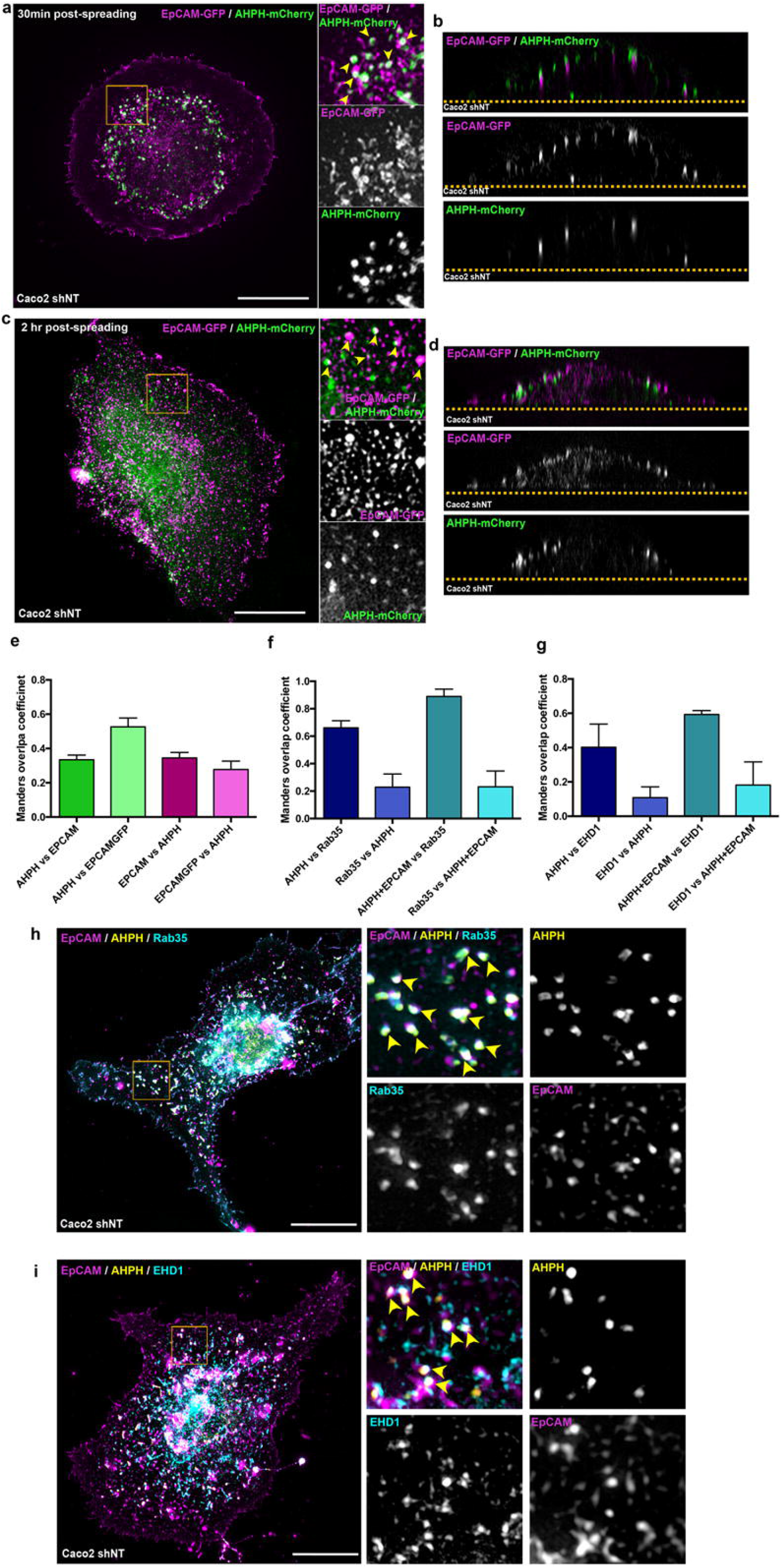
EpCAM and active RhoA co-evolve in Rab35/EHD1-positive endosomal compartments. **(a-d)** 3D-SIM microscopy analysis of EpCAM-GFP *(magenta)* and AHPH-mCherry (*green*) in unpolarized single cells during cell spreading **(a-b)** and polarity acquisition **(c-d)** in the *xy* plane **(a,c)** or the *xz* plane **(b,d)**. Areas boxed in yellow are presented on the right, where arrowheads point to colocalizations. Collagen substrate is delimited by an orange dotted line. Arrows point on colocalizations. Scale bar, 5μm**. (e)** Quantification of the Manders overlap coefficient between AHPH-mCherry versus endogenous EpCAM, AHPH-mCherry versus EpCAM-GFP, endogenous EpCAM versus AHPH-mCherry, and EpCAM-GFP versus AHPH-mCherry in polarized single Caco2 cells. (**f**) Quantification of the Manders overlap coefficient between AHPH-GFP versus Rab35-RFP, Rab35-RFP versus AHPH-GFP, AHPH-GFP+endogenousEpCAM versus Rab35-RFP, and Rab35-RFP versus AHPH-GFP +endogenousEpCAM in polarized single Caco2 cells. (**g**) Quantification of the Manders overlap coefficient between AHPH-mCherry versus EHD1-GFP, EHD1-GFP versus AHPH-mCherry, AHPH-mCherry+endogenousEpCAM versus EHD1-GFP, and EHD1-GFP versus AHPH-mCherry +endogenousEpCAM in polarized single Caco2 cells**. (h-i)** 3D-SIM microscopy analysis of EpCAM *(magenta),* AHPH-GFP *(yellow)* and Rab35-RFP or EHD1-GFP *(blue)* in polarized single cells. Areas boxed in yellow are presented on the right, where arrowheads point to colocalizations. Scale bar, 5μm.

Along this line, we evaluated the effect of EpCAM silencing on the Rab35-/EHD1-turnover of GTP-RhoA. Whereas a weak change was found at the level of Rab35-positive compartments (Figure 8a,c), the proportion of AHPH localized in EHD1-containing compartments significantly increased under EpCAM-KD (from 40% in control cells up to 60% in mutant cells; Figure 8b,d), suggesting that GTP-RhoA may remain accumulated there. To test so, we further scrutinized the dynamics of AHPH together with EHD1-positive compartments (Supplementary video 8). Tracking in control cells revealed a short residence time of AHPH in EHD1-endosomes (yellow arrowheads; Figure 8e-f; Supplementary videos 8-9). It is worth mentioning that AHPH entering EHD1-compartments exited (Figure 8e, yellow arrow; Supplementary video 9), suggesting that RhoA inactivation *per se* likely not occurs in these endosomes. However, long-residence time was observed in EpCAM-KD cells, AHPH being sequestered in EHD1-compartments (yellow arrowheads, Figure 8e-f; Supplementary videos 10 and 11). These findings showed that the cortical accumulation of the AHPH probe we observed in the absence of EpCAM reflected a block of the endosomal trafficking of GTP-RhoA-GTP during cell spreading. To provide further evidence that this endosomal pathway is required for turnover of GTP-RhoA and cell organization, we used dominant negative mutant forms of either EHD1 (mutation of glycine 65 to arginine in the P-loop domain of EHD1 which renders EHD1 cytosolic, i.e. EHD1G65R-GFP) (Naslavsky and Caplan, 2011) or Rab35 (Rab35S22N-GFP) (Kouranti et al., 2006) (Figure 8g). Overexpression of the mutant forms in control cells led to the accumulation of the AHPH location biosensor in a circular manner within the cell protrusion, nicely mimicking the effect of EpCAM silencing on RhoA-GTP distribution. Interestingly, concomitant perturbation of the epithelial cell shape was observed after EHD1G65R-GFP or Rab35S22N-GFP expression (Figure 8g). However, although transfected cells exhibit modified arrangement of actin cables and focal adhesions, the phenotype resulting from the expression of endosomal mutants differed from the one of EpCAM-KD cells (Supplementary Figure 9d), suggesting that EpCAM’s loss does not block the Rab35-EHD1 endosomal pathway process *per se* but rather the progression of GTP-RhoA there. In conclusion, our results demonstrate that active RhoA is processed along with EpCAM via the cortical endosomal road mediated by Rab35 and EHD1, and this GTP-RhoA turnover is required to ensure actin cable rearrangement and cell shape changes. Moreover, EpCAM potentiates active RhoA progression through the endosome pathway for proper turnover.

**Figure 8:**
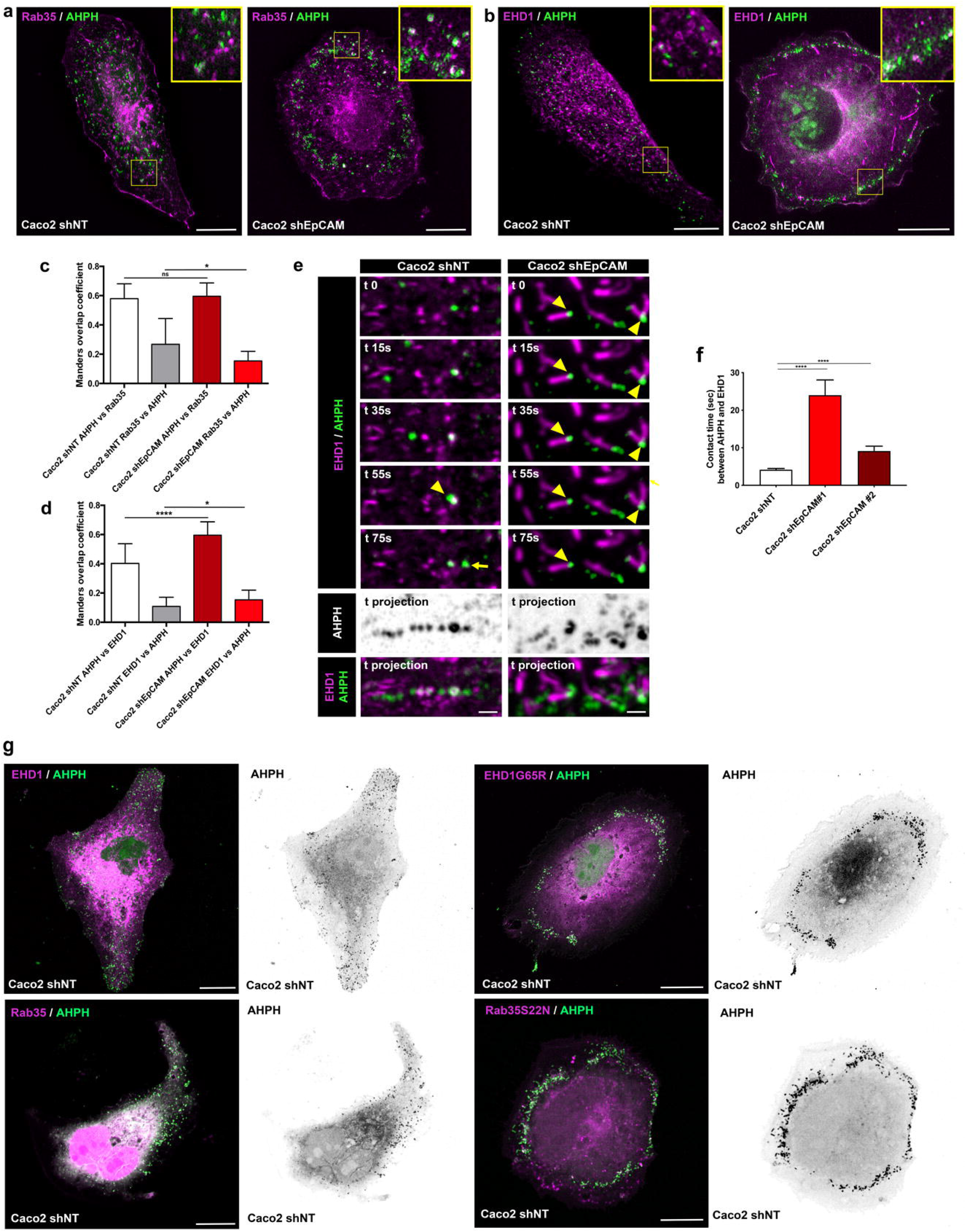
Active RhoA is blocked in Rab35/EHD1-positive endosomal compartments in the absence of EpCAM. **(a-b)** 3D-SIM microscopy analysis of the distribution of Rab35-RFP (**a**, *magenta)* or EHD1-GFP (**b**, *magenta)* together with AHPH-mCherry (*green*) in control and EpCAM-silenced cells. Areas boxed in yellow are presented on the right corner. Scale bars, 5μm. **(c)** Quantification of the Manders overlap coefficient between AHPH-GFP versus Rab35-RFP or Rab35-RFP versus AHPH-GFP in control or EpCAM-depleted cells. Unpaired t-test, \**P*=0.02. **(d)** Quantification of the Manders overlap coefficient between AHPH-mCherry versus EHD1-GFP or EHD1-GFP versus AHPH-mCherry in control or EpCAM-depleted cells. Unpaired t-test, \**P*=0.04, \*\*\*\**P* <0.0001. (**e**) Time-lapse series and maximum projection (standard deviation) of EHD1-GFP together with AHPH-mCherry in control and EpCAM-KD cells. Yellow arrowheads point at the position of colocalized AHPH and EHD1 compartments. Yellow arrow point at AHPH-positive vesicule at the exit of EHD1-postive compartment. Scale bars, 200nm. (**f**) Statistical analysis of the residence time between the AHPH probe and the EHD1-positive compartments per track in control and EpCAM-silenced cells. Measurements were made on time-lapse series of 1sec intervals. N *(Caco2 shNT)* = 8 cells (491 contacts), N *(Caco2 shEpCAM#1)* = 3 cells (55 contacts), N *(Caco2 shEpCAM #2)* = 7 cells (213 contacts). Unpaired t-test, \*\*\*\**P* <0.0001. Three independent experiments were carried out. **(g)** Confocal analysis of AHPH-mCherry (*green*) after EHD1-GFP and EHD1G65R-GFP expression *(magenta; upper pane1)* or after Rab35-GFP and Rab35S22N-RFP expression *(magenta; lower pane1)* in control Caco2 cells. Scale bars, 5μm.

## Discussion

Our study advances the understanding of contractility control during cell shaping and reveals the central participation of EpCAM in this process. Work arising from our lab and others previously pointed to EpCAM as a critical player in the spatial organization of the actomyosin network in epithelial tissues and ultimately in the apico-basal polarity and monolayer integrity (Maghzal et al., 2010, 2013; Salomon et al., 2017). EpCAM has been initially described as a “cell adhesion molecule” working at cell-cell contacts (Litvinov et al., 1994; Balzar et al., 1998). Here, we show that it is needed for isolated epithelial cells to structure their actomyosin network and properly self-organize in a front-rear axis (Figure 1e,f). We thus posit that the impact of EpCAM is a cell-autonomous general regulatory mechanism for epithelial cell plasticity. We provide further evidence of a direct implication of EpCAM in the regulation of cell contractility, where it coordinates the release of GTP-RhoA from cortical endosomes (Figures 6, 8) and may behave as a scaffolding molecule for GTP-RhoA turnover. From a mechanical point of view, what advantage would EpCAM expression give to an epithelial cell? It is interesting to note that fibroblasts or mesenchymal cells do not express EpCAM, but spontaneously self-organize and display active motility. However, in physiological conditions, EpCAM is specifically expressed in epithelial layers which, by their intrinsic nature of interfaces, are under continuous mechanical stimulation and are subjected to acute remodeling events. Furthermore, EpCAM expression often increases in epithelial tumors, that are mechanically-challenging environments (Huang et al., 2018; Keller et al., 2019; Yahyazadeh Mashhadi et al., 2019). In addition, EpCAM is widely used as a marker for detection or isolation of circulating tumor cells derived from cancers of epithelial origin such as ovarian, breast or colorectal cancers (Dementeva et al., 2017; Li et al., 2019). Here we reveal than EpCAM potentiates RhoA turnover for proper stress fiber maturation and subsequent efficient migration of individual epithelial cells. Then, EpCAM’s expression in tumor cells might maximize the cycling robustness of RhoA through its fast-endosomal turnover for proper patterning of forces generated at the cell scale, and as a consequence facilitate or sustain cancer propagation.

It is well-established that RhoA signaling dictates myosin-II-dependent contractility and stress fiber generation, and subsequently participates in cell morphogenesis and behavior (Nobes and Hall, 1995; Chrzanowska-Wodnicka and Burridge, 1996). Coordinated requirement of RhoA, Rac1 and Cdc42 activity occurs at the cell leading edge to support protrusive activity as well as rear retraction (Raftopoulou and Hall, 2004). The development and use of FRET probes showed that RhoA was actually highly activated in a 2 μm wide band at the leading edge of migrating cells and participated to protrusive activity while Rac1 and Cdc42 stabilized the protrusion for directed motion in fibroblasts (Machacek et al., 2009). Although these studies were essential to our global understanding of RhoGTPases functions, FRET analyses only provide a fixed image and low spatial resolution of their activity at a given time (Supplementary Figure 6b-d). By using a fluorescence-based location biosensor which allows the direct tracking of GTP-RhoA (Tse et al., 2012; Priya et al., 2015), our analyses reveal the transient formation of a cortical ring of active RhoA during the early steps of spreading. This RhoA zone is remodeled during late step of spreading prior to actomyosin network reorganization (Figure 6j and Figure 9). In agreement with several studies reporting the importance of proper balance of contractile forces for spreading and polarization in fibroblasts cells (Prager-Khoutorsky et al., 2011; Trichet et al., 2012), our study reveals a spatiotemporal modulation of RhoA activity during front-rear axis development. In addition, blocking of GTP-RhoA endosomal trafficking in EHD1-/Rab35-mutated or EpCAM-KD cells impairs the local regulation of RhoA signaling and subsequently the late spreading steps’ completion (Figures 6 and 8, respectively). We thus propose that a tight coupling between the remodeling of the active RhoA pool and the reshaping of actomyosin cables is necessary for correct initiation of front-rear polarity in epithelial cells.

**Figure 9:**
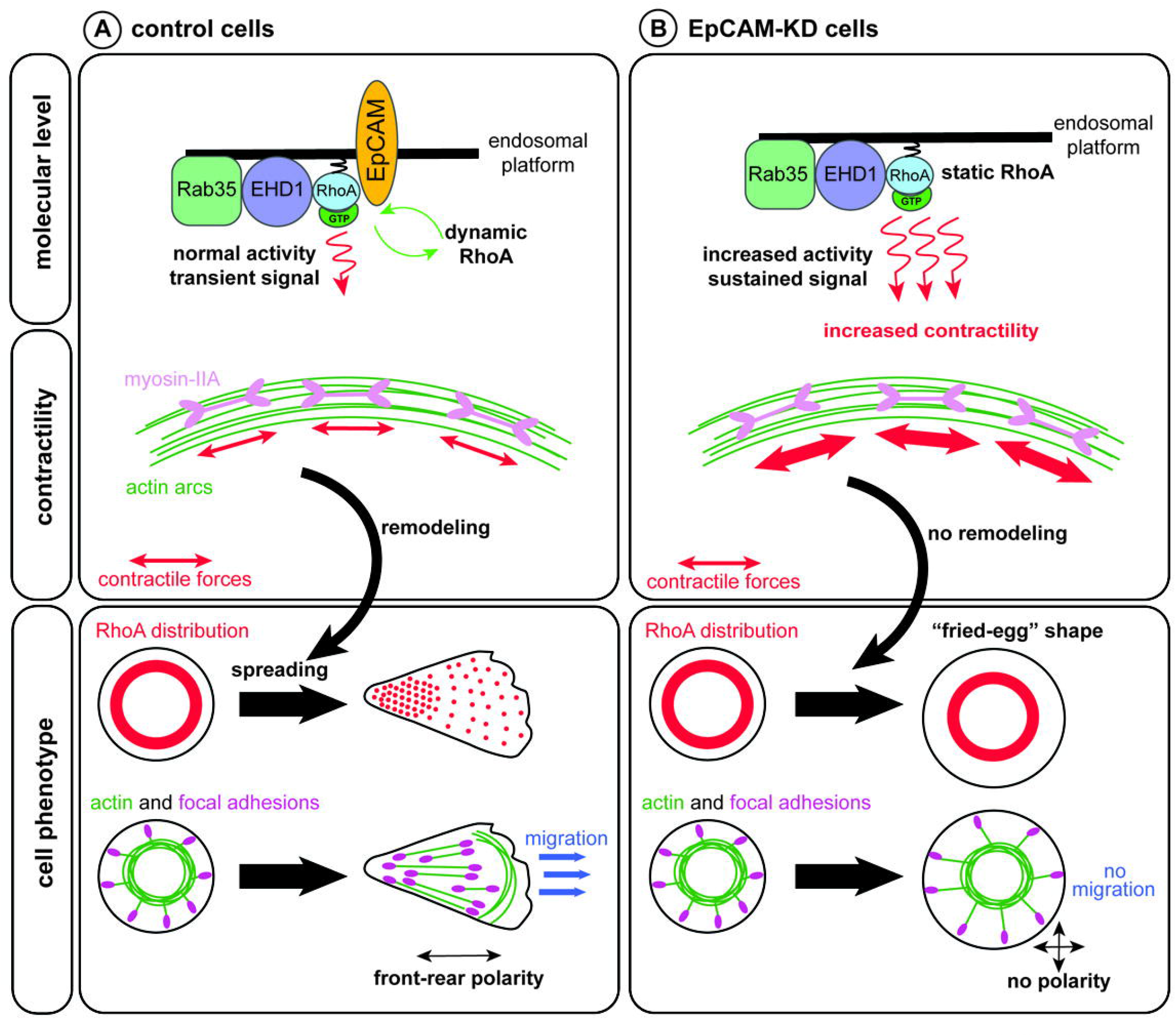
Scheme depicting the proposed model of active RhoA dynamics in control and EpCAM-KD cells during spreading. **(a)** In control cells, active RhoA (RhoA-GTP) dynamics are promoted by EpCAM, to and from the Rab35^+^/EHD1^+^ endosomal platform. The resulting transient signal induces normal Myosin-II-dependent contractility at the level of the transverse arcs during spreading. At the cellular level, dynamic RhoA-GTP can be remodeled in a front-rear gradient as the cell spreads, participating to the acquisition of front-rear polarity. Correct contractility at the levels of the transverse arcs allows the formation of ventral stress fibers, and proper actomyosin cytoskeleton reorganization to promote epithelial cell migration. **(b)** In EpCAM-KD cells, active RhoA is blocked in the endosomal platform preventing the remodeling necessary for correct spreading, symmetry-breaking and polarity establishment. RhoA sustained activity increases Myosin-II contractility at the transverse arcs level, which hinders the formation of ventral stress fibers. Active RhoA also increases formin activity, producing longer dorsal fibers in EpCAM-KD cells. The absence of active RhoA and actomyosin cytoskeleton remodeling impedes symmetry-breaking, giving EpCAM-KD cells a characteristic unpolarized “fried-egg” shape and preventing efficient cell migration.

But what drives this timing? In other words, how is this process controlled and which signal triggers GTP-RhoA exit from cortical endosomes during cell spreading? One explanation might be that a RhoA trafficking is controlled via a mechanical feedback. Previous work from Sheetz and colleagues described a sequential mechanical model of cell spreading, where each phase represents a distinct mechanical state of the cells (Dubin-Thaler et al., 2008). Whereas early cell spreading is characterized by continuous protrusive activity of the edges with very low traction forces generation (P1 phase), the P2 phase is described as a slow spreading phase during which focal adhesion form and high membrane tension occurs. Control of membrane tension appears critical for the progression through this P2 phase (Gauthier et al., 2012), and would be ensured by coordinated regulation of membrane trafficking events and myosin-II contractility (Apodaca, 2002; Gauthier et al., 2011; Pontes et al., 2017). For instance, inhibition of myosin-IIA activity keeps fibroblasts blocked in P2 phase (Cai et al., 2010), and reduction of membrane tension triggers endocytosis pathway (Thottacherry et al., 2018). Our findings show that EpCAM-KD cells fail to complete P2 phase and display isotropic and, to some extent, C-shapes (Figure 1d-i; Supplementary Figure 1), as do myosin-IIA mutant fibroblasts (Cai et al., 2010). There, loss of GTP-RhoA turnover leads to persistent RhoA signaling at the cell cortex, causing continuous actomyosin contractility in the dorsal domain. Consequently, protrusion stiffness remains very high in EpCAM-KD cells (Figure 3a-c), suggesting an increase of membrane tension. Along this line, low dose treatment with Y-27632 or blebbistatin, which partially restores cell organization of EpCAM-KD cells (Figure 5e-f), may lead to a softer cortex in the protrusion, more prompt to cell deformation. We thus hypothesize that active RhoA zone remodeling would take place during the P1-P2 transition phase and/or at the onset of P2 phase. The dynamic mechanism of RhoA signaling described herein would support this earlier spreading model, and we propose that the exit of RhoA from endosomal compartment may constitute a spatio-temporal signal in this sequence. The RhoA trafficking may provide a rapid response to high membrane tension, and thus contribute to the progression through the late spreading cycle. Another explanation would involve actomyosin activity by itself. Several studies reported that pulsatile contractions take place at the medio-apical and junctional pools of actomyosin in diverse species to facilitate cell shapes changes and tissue morphogenesis (Mason et al., 2016; Wu et al., 2014). Constitutively active myosin-II phospho-mutant expression disrupts pulsatile contractility and delays tissue invagination during *Drosophila* gastrulation (Vasquez et al., 2014). Similarly, we could propose that persistence of GTP-RhoA at the cell cortex would hinder efficient contractility to ensure the transition from radial and circumferential cables to stress fibers.

In contrast to the commonly accepted view of RhoA cycling at the plasma membrane, our work places a large pool of active RhoA in trafficking pathways, leading to the following question: why the need of a GTP-RhoA endosomal turnover? Several hypotheses could be envisioned. It may allow fine tuning of GTP-RhoA distribution during spreading and migration in an ever-changing cell shape environment. Moreover, contractility pulses may be related to cyclic activation / inactivation of RhoA and may thus potentiate the efficiency of contractility (Teo and Yap, 2016). The intracellular trafficking of RhoA might also be used to target GTP-RhoA to other intracellular or plasma membrane domains to meet regulators or effectors. RhoA cycling between its active and inactive state is promoted by the sequential action of its guanine nucleotide exchange factors (GEFs) that promote GTP-loading, and GTPase-activating proteins (GAPs) that favor GTP hydrolysis (Rossman et al., 2005; Bos et al., 2007; Hodge and Ridley, 2016). Recent works and reviews pointing towards the same surprising distributions of active GTPases proposed that the patterns of RhoA activity relied on the subcellular distribution of GEFs and GAPs in function-oriented domains, rather than on the GTPase’s location in itself (Fritz and Pertz, 2016). More than 80 RhoGEFs and GAPs are reported in the human genome (Lawson and Ridley, 2018), suggesting that the RhoGTPases cycling is more specifically regulated than a simple ON-OFF switch. Depending on the trafficking pathway used to remodel the RhoA zone, different combinations of GEFs and GAPs would grant an extreme precision for the spatio-temporal control of RhoA activity. The careful characterization of subcellular patterning of GEFs and GAPs might be laborious but still remains an important subject of study for the future. An alternative view is that endosomal trafficking may regulate RhoA signaling by removing GTP-RhoA away from the actomyosin cables region and/or to prevent the gathering of GTP-RhoA with its effectors ROCK1, mDia and MLCP.

We propose a model where EpCAM-mediated endosomal remodeling allows the local modulation of RhoA signaling in space and time during epithelial cell spreading (Figure 9). The canonical picture states that activated RhoA is restricted at plasma membrane (Garcia-Mata et al., 2011; Hodge and Ridley, 2016). While transmembrane receptor trafficking, such as for integrins or cadherins, has been under intense scrutiny in the context of cell migration and cancer metastasis (Mellman and Yarden, 2013; Paul et al., 2015; Kajiho et al., 2018), the link between traffic and Rho GTPases is only now coming to the fore and our data support the new idea that “endosomes serve as a hub for Rho GTPase activation and spatiotemporal signal generation” (Phuyal and Farhan, 2019). Recent studies highlighted the presence of active Rho GTPases in intracellular membranes and argue for an origin of Rho signaling from the endosomal network (Phuyal and Farhan, 2019). Vassilev et al. recently showed that in neural crest cells RhoA trafficking is mediated by Rab11-positive recycling pathway (Vassilev et al., 2017). However, our findings revealed that, even though a small proportion of active RhoA was indeed carried by Rab11-positive compartments (Supplementary Figure 9b-c), the major endosomal pathway for RhoA is in fact mediated by Rab35/EHD1-positive compartments in epithelial cells (Figure 7). In cortical endosomes, Rab35 and EHD1 control a fast recycling process that parallels the canonical Rab11 pathway (Klinkert and Echard, 2016). Whereas Rab35 functions during early step of endosomal recycling, EHD1 is required for late endosomal scission events (Kouranti et al., 2006; Klinkert and Echard, 2016; Cauvin et al., 2016). Even though Rab35 functions’ in cell migration and adhesion vary according to the cell types or migration assays, it emerges as a pivotal component during cancer progression for receptor presentation, actin dynamics and cell polarity (Shaughnessy and Echard, 2018; Corallino et al., 2018). A link between Rab35 and RhoA was mentioned previously although not clearly demonstrated nor with any apparent impact (Chevallier et al., 2009). At this stage, we can only speculate that distinct endosomal compartments would constitute cortical reservoirs of GTP-RhoA that cells might differentially use to reorganize the contractile network and force generation in response to diverse situations or external cues. We realize that, for now, our study only provides a small window on the spatio-temporal regulation of contractility in epithelial cells, and a complete view of the upstream regulatory mechanisms and trafficking pathways involved in the regulation of RhoA activity will deserve future in-depth analyses. Moreover, as several reports pointed out the importance of RhoA activity for the maintenance of myosin-II activity and junctional integrity (Arnold et al., 2017; Priya et al., 2017), whether a similar endocytic mechanism occurs at cell contacts should be tested in the future.

In summary, our results unveil endosomal trafficking as a key mechanism of spatial-temporal control of RhoA during stress fiber formation and cell polarity acquisition in epithelial cells, and provide a mechanistic understanding with the characterization of EpCAM-mediated active RhoA turnover through the cortical endosomes.

## Materials & Methods

### Cell culture

Caco2, U2OS and HeLa cells were kindly provided by Dr. S. Robine (Curie Institute, Paris) and Valérie Doye (Institut Jacques Monod, Paris), respectively. Caco2 and MDCK cells were routinely grown in DMEM 4.5 g/l glucose supplemented with 20% (Caco2 cells) or 10% fetal bovine serum and 1% penicillin-streptomycin (Gibco, Thermo Fischer Scientific, Waltham, MA, USA) for maximum 9 passages. The culture medium was renewed every 2-days. For all experiments cells were plated on collagen-coated substrates, obtained by adsorption of collagen I (Sigma) that was incubated at room temperature for 1h at 100 μg/mL in 0.02N acetic acid, and washed with PBS before cell seeding.

EpCAM reduction was carried out by lentiviral delivery of shRNA constructs directed against human *EPCAM shEpCAM#1:* TRCN0000073734 5’-CCGGGCCGTAAACTGCTTTGTGAATCTCGAGATTCACAAAGCAGTTTACGGCTTTTTG-3’, and *shEpCAM#2:* TRCN0000073737 5’-CCGGCGCGTTATCAACTGGATCCAACTCGAGTTGGATCCAGTTGATAACGCGTTTTTG-3’ designed and cloned into the lentiviral pLKO.1 puromycin resistant vector Mission shRNA lentiviral Transduction particle (Sigma Aldrich). Control Caco2 clones *(shNT)* were generated using pLKO.1-puro non-target shRNA control transduction particles SHC016V (5’-CCGGGCGCGATAGCGCTAATAATTTCTCGAGAAATTATTAGCGCTATCGCGCTTTTT-3’). shRNA-resistant EpCAM sequence (shEpCAM#1 resistant: 5’-ATGGCGCCCCCGCAGGTCCTCGCGTTCGGGCTTCTGCTTGCCGCGGCGACGGCGACTTTTGCCGCAGCTCAGGAAGAATGTGTCTGTGAAAACTACAAGCTGGCTGTGAATTGTTTCGTCAACAATAATCGTCAATGCCAGTGTACTTCAGTTGGTGCACAAAATACTGTCATTTGCTCAAAGCTGGCTGCCAAATGTTTGGTGATGAAGGCAGAAATGAATGGCTCAAAACTTGGGAGAAGAGCAAAACCTGAAGGGGCCCTCCAGAACAATGATGGGCTTTATGATCCTGACTGCGATGAGAGCGGGCTCTTTAAGGCCAAGCAGTGCAACGGCACCTCCATGTGCTGGTGTGTGAACACTGCTGGGGTCAGAAGAACAGACAAGGACACTGAAATAACCTGCTCTGAGCGAGTGAGAACCTACTGGATCATCATTGAACTAAAACACAAAGCAAGAGAAAAACCTTATGATAGTAAAAGTTTGCGGACTGCACTTCAGAAGGAGATCACAACGCGTTATCAACTGGATCCAAAATTTATCACGAGTATTTTGTATGAGAATAATGTTATCACTATTGATCTGGTTCAAAATTCTTCTCAAAAAACTCAGAATGATGTGGACATAGCTGATGTGGCTTATTATTTTGAAAAAGATGTTAAAGGTGAATCCTTGTTTCATTCTAAGAAAATGGACCTGACAGTAAATGGGGAACAACTGGATCTGGATCCTGGTCAAACTTTAATTTATTATGTTGATGAAAAAGCACCTGAATTCTCAATGCAGGGTCTAAAAGCTGGTGTTATTGCTGTTATTGTGGTTGTGGTGATAGCAGTTGTTGCTGGAATTGTTGTGCTGGTTATTTCCAGAAAGAAGAGAATGGCAAAGTATGAGAAGGCTGAGATAAAGGAGATGGGTGAGATGCATAGGGAACTCAATGCATAA-3’) was provided by Invitrogen and cloned into a pEGFP-N 1 backbone using the following primers (Eurofins genomics): Forward: 5’-aattctgcagtcgacggtaccATGGCGCCCCCGCAGGTC-3’, Reverse: 5’-caccatggtggcgaccaggtggatcccgggT GCATT GAGTTCCCT AT GCATCTCA-3’. shEpCAM#1-resistant Caco-2 clones were generated by transfection with Lipofectamine 2000 (Thermo Fisher Scientific) according to the manufacturer’s instructions, and selection was perfomed in DMEM supplemented with 20% FBS, 10% penicillin/streptomycin, 2 μg/ml puromycin and 0.5 mg/ml geneticin (Life Technologies, Paisley, UK).

Plasmid and siRNA transient transfections were performed using Lipofectamine 2000. ACTN4 silencing was carried out using two siRNA targeting human ACTN4 mRNA, from Sigma Aldrich. siRNA ACTN4 #1: 5’-CUUCUCUGGUGCCAGAGAA[dT][dT]-3’, siRNA ACTN4 #2: 5’-GACAUGUUCAUCGUCCAUA[dT][dT] – 3’. EpCAM reduction in MDCK cells was carried out using two siRNA targeting dog EpCAM mRNA, purchased from Invitrogen: siRNA EpCAM #1: 5’-UUCAUAACCAAACAUUUGGUUGCCA-3’, siRNA EpCAM #2: 5’-UGAUUGAGAGCUGCCUUUCUAUUUA-3’. EpCAM-GFP was purchased from Origene (NM_002354, CAT# RG201989). Rab11-dominant negative mutant was purchased from Addgene (# 12678). AHPH-GFP, AHPH-mCherry, AHPH^A740D^, ROCK1-GBD-GFP and mDia-GBD-GFP constructs were a kind gift from Dr. A.Yap (Brisbane University, Australia). GFP-Rho wt was a gift from Dr Anne Blangy (CRBM, Montpellier, France). Rab5a-mCherry and Rab11a-GFP constructs were a gift from Dr. C. Wunder (Curie Institute, France). EHD1-GFP and EHD1-G65R-GFP were a gift from Dr S. Caplan (University of Nebraska Medical Center, NE, USA). Rab35-RFP and Rab35-S22N-RFP were provided by Dr A. Echard (Pasteur Institute, Paris, France). LifeAct-GFP was from Addgene.

### Antibodies and reagents

Rabbit polyclonal antibody directed against EpCAM (#ab71916, IF dilution, 1:100) and mouse monoclonal antibody directed against zyxin (#ab58210, IF dilution 1:100) were from Abcam. Mouse monoclonal antibodies directed against paxillin (# 5H11, IF dilution, 1:100) and talin (#TA205, IF dilution, 1:100) were from Merck Millipore. Rat monoclonal antibody against activated β1-integrin was from BD Biosciences (#, IF dilution: 1/100). Rabbit polyclonal antibody against vinculin (# V4139, IF dilution, 1:100) was from Sigma-Aldrich. Rabbit polyclonal antibody directed against MLC2 (#3672, WB dilution 1:1,000) and P-MLC2 (T18/S19, #3674S IF dilution 1:100, WB dilution 1:500) were from Cell Signaling (Danvers, MA, USA). Rabbit monoclonal antibody directed against α4-actinin was from Life Technologies (#42-1400, IF dilution, 1:100). Monoclonal antibody directed against GAPDH (#60004-1-Ig, clone 1E6D9, WB dilution, 1:500) was from Proteintech (Chicago, IL, USA). Rabbit polyclonal antibody directed against Myosin-IIA (#909801, IF dilution 1:100) was from Biolegend (Princeton, NJ, USA). Mouse monoclonal antibody directed against α1-actinin (#TA500072S, IF dilution 1:100) was from Origene. Phalloidin-Alexa488, 568 or 647 were from Life Technologies. Blebbistatin, Y-27632, CK666, ML-7, SMIFH2 were from Sigma Aldrich (SaintLouis, MO, USA). NSC-23766 was from Tocris (Bio-Techne, France), and CN03 was from Cytoskeleton (Denver, CO, USA).

### Biochemical analysis

For Western blot, cell lysates were prepared 1 or 21 days after plating, for protein detection in single cells or polarized monolayer respectively. Cells were lysed for 30 min using the following lysis buffer: 50 mM Tris/HCl pH 8.0, 150 mM NaCl, 1 mM DTT, 0.5% NP-40, 1% Triton X-100, 1 mM EGTA, 1mM EDTA, with complete protease inhibitor cocktail and phosphatase inhibitor PhosSTOP (Roche, Basel, Switzerland). Insoluble debris were removed by centrifugation at 13,000 g for 15 min. Total protein content was measured by Bradford assay (Biorad). For each condition, 50mg of proteins were loaded per well in Novex Tris-Glycine pre-cast gels (Thermo Fischer Scientific)) and transferred on nitrocellulose membranes using iBlot Dry blotting system (Thermo Fischer Scientific)). Proteins were detected with either HRP-linked goat anti-mouse IgG antibody (dilution 1:10,000; Sigma-Aldrich) or HRP-linked donkey anti-rabbit IgG antibody (dilution 1:10,000, GE Healthcare, Buckinghamshire, UK), and SuperSigna West Femto Maximum Sensitivity Substrate (Thermo Fischer Scientific), and visualized on ImageQuant LAS4000 (GE-Healthcare). Signal quantification was performed using Fiji software.

For immunoprecipitation, cells were lysed as described above. Lysates were precleared with protein A-Sepharose beads (Sigma-Aldrich) for 1 h, incubated with antibodies overnight at 4°C, and incubated with newly prepared protein A–Sepharose beads the next day for 2 h. The beads were washed three times with the lysis buffer. Precipitates were separated by SDS-PAGE and analyzed by immunoblotting.

### Immunostaining

Cells were fixed using 4% paraformaldehyde for 15 min, then permeabilized using 0.02% saponin solution in PBS for 20 min. 0.02% saponin/1% BSA solution was used for a 30min blocking step, before proceeding to incubation with the primary antibody at 4°C overnight. The next day, secondary antibody was added after 3 washing steps in PBS, and left to incubate for 2h at RT. Except for SIM analysis where Vectashield medium was used, all staining were mounted in Mowiol.

### Live imaging

Live cell spreading and migratory assays were performed with the Biostation (Nikon, Tokyo, Japon) using the 20x objective. Time-lapse images were taken every 10 min for 2 to 4 hours. Cell were treated with mitomycin C (10 μg/mL) (Sigma-Aldrich) for 1h to prevent division, before seeding on collagen-coated glass bottom fluorodishes (#FD35-100, World Precision Instruments, Sarasota, FL, USA). Cell tracks measurements and graphs were obtained using MATLAB (Mathworks, Natick, MA, USA).

Cell edge protrusive activity was analyzed using kymographs from 10 min time-lapse movies with 5s frame rate, obtained on a wide field DMI6000 microscope (Leica, Wetzlar, Germany) using x100/1.4NA Plan apochromatic oil objective. Briefly, four lines per cell normal to a free cell edge were used to generate kymographs from which protrusion and retraction rates, time of protrusion and retraction, amplitude and period were measured with ImageJ.

Actin dynamics experiments were performed using an inverted DMI8 Leica microscope equipped with a CSU-W1 spinning disk head (Yokogawa – Andor), using a x100 1.4 NA oil objective. Images were acquired every 5 or 12min for 2-4h. Active RhoA and EHD1 dynamics were followed on the same microscope, for 2min with a 1s frame rate. For the analysis of AHPH probe and EHD1-positive compartment contact time, AHPH-mCherry vesicles were manually tracked and EHD1-GFP contact length was manually assessed.

Color-coded t-projection of AHPH-mCherry were generated from spinning disc acquisitions, as described by O’Neill and colleagues (Bach et al., 2014). 10 frame-t-stack were selected from timelapse series. The first image (*t0*) was false-colored green, the last image (*t9*) was false-colored in blue, and the intervening time points (*t2-8*) were submitted to t-projection and shown in red (*t projection*), and images were merged.

### Structured Illumination Microscopy

3D-Structured Illumination Microscopy (SIM) was performed on a Zeiss Elyra Microscope coupled to an optovar 1.6, 63X objective and a camera EM CCD Andor SIM. During z-stack acquisition, 5 rotations were applied. Deconvoluted structured illumination images were generated by Zen software, and images were merged in ImageJ.

### Atomic force microscopy

Cells were cultured in DMEM, 20% FBS, 1× penicillin-streptomycin, and 2 μg/ml puromycin. Plastic petri dishes (TPP, Switzerland) were incubated in 100 μg/mL rat tail collagen I (Gibco A10483-01) in 0.1% acetic acid at 4°C overnight on a 60 rpm shaker. Cells were seeded at low density and allowed to adhere at least 5 hours. AFM nanoindentation experiments were performed with a Nanowizard 4 (JPK Instruments, Germany) in QI™ mode. The imaging buffer was Leibovitz L-15 medium supplemented with 20% FBS and 1× penicillin-streptomycin and experiments were performed at 37°C with a petri dish heater. PFQNM-LC-A-CAL cantilevers (Bruker, USA) were used; the nominal tip radius is 70 nm, the semi-vertical angle is 17°, the probe spring constant was provided by the manufacturer, and the optical lever sensitivity was determined by the thermal tuning method. Force-indentation curves were collected with 100 μm/s probe velocity, 400 pN trigger force, and variable indentation-retraction distance (scanning frequency) over a 60 μm^2^ area with 128×128-pixel resolution (each cell scan lasted ~10 min, the fast axis was horizontal in images shown). Data was analyzed using a custom-built MATLAB program; Young’s modulus values are fit along the entire force-indentation curve using a linearization scheme (Staunton et al., 2016) with the Hertz model modified for a thin sample adhered to an infinitely rigid substrate (Garcia and Garcia, 2018). The height at each pixel was determined from the contact point and the Poisson’s ratio was assumed to be 0.5.

### Traction Force Microscopy

Soft polydimethylsiloxane (PDMS) substrates of 15kPa rigidity containing red 200nm fluorescent beads (Life technologies) were prepared by mixing CY52-276 kit components (Dow Corning Toray) at 1:1 ratio and letting it spread and cure on glass bottom fluorodishes overnight. Collagen I (Sigma Aldrich) was adsorbed on the surface on the substrate as described above. Cells were plated on the substrate and imaged for 22-24h, at a 6 min interval in the Biostation using the x20 objective. Images were aligned to compensate for experimental drift before analysis of the beads displacement was performed, using Particle Imaging Velocimetry (PIV) script in MATLAB. From the displacement data, Fourrier transform traction cytometry (FTTC) plugin in ImageJ (available at https://sites.google.com/site/qingzongtseng/tfm) was used to estimate the traction forces exerted on the substrate. The total traction force exerted by cells was calculated by summing the magnitudes of the traction vectors under and near the cell of interest and multiplying by the area covered by those vectors.

### FRET analyses

The FRET probe Raichu-1502 (also named Raichu-RBD (Yoshizaki et al., 2003)) was transfected into Caco2 cells 48h before the experiment. Spectral imaging was performed on a confocal LSM780 microscope (Zeiss, Zen software) with x63/1.4NA plan apochromat oil-immersion objective. CFP was excited by the 458-nm laser line of an Argon laser and emission was sampled at a spectral resolution of 9-nm within a 444–570-nm range. ImageJ was used to process images for analyses. FRET ratio was calculated as the ratio between the YFP and CFP signal.

### Statistical analysis

All statistical analyses were performed using Prism (GraphPad Software, San Diego, CA, USA, version 7.0). Unless otherwise stated, experiments were replicated 3 times independently and comparison between samples were done without Gaussian distribution assumption of the data, meaning comparisons were carried out using Mann-Whitney test for two conditions comparison, or Kruzkal-Wallis test and Dunn’s multiple comparison tests for three and more conditions. P-values met the following criteria * p < 0.05, ** p < 0.01 and *** p < 0.001.

Triple colocalization was analyzed from confocal stacks using a MATLAB-based custom program. Briefly, fluorescence was first segmented in each channel using local thresholding (Phansalkar method; (Neerad Phansalkar et al., 2011) and a local 2-D median filter with user-defined 16 neighborhood size was applied to remove noise. Colocalization was then measured using the Manders split coefficients M1 and M2 (Manders et al., 1993) as such: 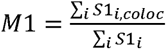 and 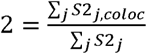, where S1_i,coloc_ = S1_i_ if S1_i_ > 0 AND S2_i_ > 0, and S2_j,coloc_ = S2_j_ if S2_j_ > 0 AND; S1_j_ > 0. The Matlab Code or the standalone user interface can be shared upon request.

## Data availability

The data that support the findings of this study are available from the corresponding author upon request. Raw western blots are already presented in Supplementary Figures 10 and 11.

## Acknowledgments

We thank Alpha Yap, Arnaud Echard and René-Marc Mège for helpful discussions. We thank Sarah Benlamara, Arthur Radoux and Dylan Gourmelin for technical help. Confocal microscopy and high-resolution 3D-SIM analyses were performed in the ImagoSeine microscopy facility (Institut Jacques Monod, IJM). This work was supported by grants from the Groupama Foundation – Research Prize for Rare Diseases 2017 (to D.D), the Fondation pour la Recherche Médicale (FRM, #FDT201805005185) (to C.G.), the Association Prolific (prix « Graine de Chercheur » 2017) (to C.G.), the LabEx “Who Am I?” #ANR-11-LABX-0071 and the Université de Paris IdEx #ANR-18-IDEX-0001 funded by the French Government through its “Investments for the Future” program (to D. D), and the Human Frontier Science Program (RGP0038/2018) (to C.T.L. and D.D).

## Abbreviations

CTE: Congenital Tufting Enteropathy;
EpCAM: epithelial-cell adhesion molecule;
FA: focal adhesion;
IF: immunofluorescence;
KO: knock-out;
MLC2: myosin light chain 2;
P-MLC2: phosphorylated myosin light chain 2;
SIM: structured illumination microscopy;
TA: transverse arcs;
WB: Western blot.

## Supplementary Information

(Supplementary Figures 1 to 9, Supplementary Videos 1 to 11, and original western blots used in the manuscript displayed in Supplementary Figures 10 and 11) is available in the online version of the paper.

## Author contributions

C.G., B.D., P.M., E.G., S.B. and D.D. performed the experiments. C.G., and D.D. designed the experiments. C.G., S.D.B., B.D., P.M., J.D.A., E.G. and D.D. performed analyses. C.G., B.L. and D.D. coordinated the overall research and experiments, and wrote the manuscript.

## Conflict of interest

The authors declare no conflict of interest.

## Materials and correspondence

Correspondence and material requests should be addressed to D. Delacour.

## Supplementary Figure Legends

**Supplementary Figure 1:** Phase contrast time-lapse of EpCAM-silenced cells during cell spreading showing C-shape development. Yellow arrows point on symmetry breaking events Scale bar, 5μm.

**Supplementary Figure 2: (a)** Confocal analysis of the distribution of α4-actinin *(magenta)* and myosin-IIA (*green*) in control and EpCAM-depleted cells. (**b**) Confocal analysis of the distribution of ß1-integrin, talin, vinculin and zyxin in control and EpCAM-silenced cells. Accumulated z-stack are presented. Scale bar, 5μm. **(c**) Analysis of the distribution of focal adhesions according to their size in control *(Caco2 shNT)* and EpCAM-depleted *(Caco2 shEpCAM#1 and #2)* cells. N *(Caco2 shNT)* = 31 cells, N *(Caco2 shEpCAM#1)* = 30 cells, N *(Caco2 shEpCAM #2)* = 30 cells. (**d**) Analysis of the mean size of focal adhesions in control *(Caco2 shNT)* and EpCAM-depleted *(Caco2 shEpCAM#1 and #2)* cells. Kruskal-Wallis test, *P* <0.0001. Values are mean±SD. N *(Caco2 shNT)* = 31 cells, N *(Caco2 shEpCAM#1)* = 30 cells, N *(Caco2 shEpCAM #2)* = 30 cells. For each experiment, three independent experiments were carried out.

**Supplementary Figure 3:** (**a**) Confocal analysis of actin *(gray)* and paxillin *(red)* in Caco2 shEpCAM cells after EpCAM rescue through the expression of an EpCAM-GFP construct resistant to the shRNA (EpCAMr-GFP, *green*). Accumulated z-stack are presented. Scale bar, 5μm. **(b)** Confocal analysis of actin (*green*) and paxillin *(magenta)* distribution in control and EpCAM-depleted cell islands. Accumulated z-stack are presented. Scale bar, 5μm.

**Supplementary Figure 4: (a)** Western blot analysis of EpCAM expression in control or EpCAM siRNA-treated MDCK cells. α-tubulin was used as a loading control. **(b)** Statistical analysis of EpCAM expression in control and siRNA-treated MDCK cells. (extinction of 59% and 32% for siRNA #1 and #2 respectively). Three independent experiments were carried out.

**Supplementary Figure 5: (a)** Western blot detection of α4-actinin (*upper panel*) and EpCAM *(lower panel)* after immunoprecipitation of α4-actinin or EpCAM from Caco2 shNT cell extracts. **(b)** Western blot analysis of α4-actinin expression in control *(Luciferase siRNA)* or α4-actinin-deprived *(α4-actinin siRNA)* Caco2 shNT and Caco2 shEpCAM cells. GAPDH was used as loading control. **(c)** Confocal analysis of α4-actinin *(magenta),* actin (*green*) and paxillin *(red)* distribution in α4-actinin-depleted Caco2shNT cells *(upper panel)* or in α4-actinin-depleted Caco2 shEpCAM cells *(lower panel).* Accumulated z-stack are presented. Scale bar, 10μm.

**Supplementary Figure 6:** (a) Confocal analysis of paxillin *(magenta)* and actin (*green*) in EpCAM-depleted cells upon DMSO, blebbistatin 2 μM, blebbistatin 10 μM, Y-27632 10 μM treatment for 1 hour. Accumulated z-stack are presented. Scale bars, 5μm. (**b**) Scheme presenting the principle of the FRET probe to measure RhoA activity developed by Matsuda and colleagues (Yoshizaki et al., 2003). (**c**) FRET intensity maps were generated in control and EpCAM-depleted cells with LUT table “16 colors” from ImageJ. Scale bars, 10μm. (**d**) Statistical analyses of FRET intensity in control and EpCAM-depleted Caco2 cells. FRET ratio in Caco2 shNT cells = 1.52±0.24, Caco2 shEpCAM cells# 1 = 1.06±0.49 and Caco2 shEpCAM cells#2 = 0.96±0.42 (mean±SD). One-way Anova test, \**P*=0.04. Two independent experiments were carried out. (**e**) Confocal analysis of paxillin *(magenta)* and actin (*green*) in EpCAM-depleted cells upon SMIFH2 2 nM, ML-7 10 μM, CK666 50 μM or NSC23766 50 μM treatment for 1 hour. Accumulated z-stack are presented. Scale bars, 5μm.

**Supplementary Figure 7:** Confocal analysis of paxillin *(magenta)* and actin (*gray*) in control and EpCAM-depleted cells transfected with RhoA-GFP, RhoA V14-GFP or RhoA N19-GFP constructs (*green*). Accumulated z-stack are presented. Scale bars, 5μm.

**Supplementary Figure 8: (a-c)** Confocal analysis of total RhoA (Rho-GFP, *green)* **(a)**, RhoA-GTP binding domain of ROCK1 (ROCK1-GBD-GFP, *green)* **(b)** or RhoA-GTP binding domain of mDia (mDia-GBD-GFP, *green*), together with GTP-RhoA (AHPH-mCherry, *magenta)* in control Caco2 cells. (**d**) Confocal analysis of the localization of the wt AHPH-GFP and the mutant form AHPH^A740D^-GFP in control Caco2 cells. Scale bar, 5 μm.

**Supplementary Figure 9: (a)** Confocal analysis of AHPH-mCherry in Caco2 shEpCAM cells after rescue with EpCAMr-GFP (*green*) transfection. Areas boxed in yellow are presented on the right. Scale bar, 5 μm. **(b)** Confocal analysis of AHPH-GFP (*green*) and Rab11-mCherry *(magenta)* in control Caco2 cells. Scale bar, 5μm. **(c)** Quantification of the Manders overlap coefficient between AHPH-GFP versus Rab11-mCherry in Caco2 cells. (**d**) Confocal analysis of actin (*gray*) and paxillin *(magenta)* distribution after EHD1G65R-GFP transfection in control Caco2 cells. Scale bar, 5μm.

**Supplementary Figures 10 and 11:** Original western blots used in the manuscript.

## Supplementary Video Legends

**Supplementary Video 1:** 3-hour time lapse imaging of Caco2 shNT cells during spreading and polarity acquisition. Images were acquired every 6 min. Frame rate is 15fps.

**Supplementary Video 2:** 3-hour time lapse imaging of Caco2 shEpCAM cells during spreading. Images were acquired every 6 min. Frame rate is 15fps.

**Supplementary Video 3:** 4-hour time lapse imaging of Caco2 shEpCAM cells during spreading, showing symmetry breaking events and C-shape acquisition. Images were acquired every 6 min. Frame rate is 15fps.

**Supplementary Video 4:** 2-hour time lapse spinning-disc acquisition of Lifeact-GFP dynamics *(gray)* in Caco2 shNT cells. Images were acquired every 5min. Frame rate is 10fps.

**Supplementary Video 5:** 2-hour time lapse spinning-disc acquisition of Lifeact-GFP dynamics (*gray*) in Caco2 shEpCAM cells. Images were acquired every 5min. Frame rate is 10fps.

**Supplementary Video 6:** 2-min time lapse spinning-disc acquisition of AHPH-mCherry dynamics (*gray*) in Caco2 shNT cells. Images were acquired every 5sec. Frame rate is 10fps.

**Supplementary Video 7:** 2-min time lapse spinning-disc acquisition of AHPH-mCherry dynamics (*gray*) in Caco2 shEpCAM cells. Images were acquired every 5sec. Frame rate is 10fps.

**Supplementary Video 8:** 1-min time lapse spinning-disc acquisition of AHPH-mCherry dynamics *(red)* together with EHD1-GFP *(green)* in Caco2 shNT cells. Images were acquired every 5sec. Frame rate is 15fps.

**Supplementary Video 9:** Close-up of AHPH-mCherry dynamics *(red)* together with EHD1-GFP (*green*) during 15-sec time lapse spinning-disc acquisition in Caco2 shNT cells. Images were acquired every 5sec. Frame rate is 10fps.

**Supplementary Video 10**: 1-min time lapse spinning-disc acquisition of AHPH-mCherry dynamics *(red)* together with EHD1-GFP (*green*) in Caco2 shEpCAM cells. Images were acquired every 5sec. Frame rate is 15fps.

**Supplementary Video 11**: Close-up of AHPH-mCherry dynamics *(red)* together with EHD1-GFP (*green*) during 30-sec time lapse spinning-disc acquisition in Caco2 shEpCAM cells. Images were acquired every 5sec. Frame rate is 10fps.

## Notes

### Competing Interest Statement

The authors have declared no competing interest.

